# Epistatic mutations in ISC metabolism synergize with cell cycle regulation and the PPP to enhance xylose fermentation and acetic acid tolerance in industrial yeast

**DOI:** 10.1101/2025.08.04.668411

**Authors:** Leandro Vieira dos Santos, Gisele Cristina de Lima Palermo, Paulo Emílio dos Santos Costa, Ludimila Dias Almeida, Marcelo Falsarella Carazzolle, Gonçalo Amarante Guimarães Pereira

## Abstract

Efficient utilization of complex biomass-derived sugars and tolerance to inhibitors are key requirements for the viability of lignocellulosic-based biorefineries. In this study, a two-stage evolution of an industrial yeast strain engineered with a xylose isomerase pathway yielded strain AceY.14, which exhibited improved fermentative performance and increased tolerance to acetic acid. Whole-genome sequencing of the evolved strain identified SNPs in *ZWF1*, a component of the pentose phosphate pathway (PPP), and in the G1 cyclin gene *CLN3*, both of which were functionally validated through CRISPR and reverse engineering. The *zwf1^E191D^* mutation reduced xylitol accumulation, alleviating inhibition of xylose isomerase and enhancing flux through the non-oxidative branch of the PPP, while the frameshift *cln3^T556fs^*mutation unexpectedly improved acetic acid tolerance and xylose consumption in the evolved strain, also affecting cell size and growth. Genome sequencing of AceY.14 also revealed a significant reduction in the *xylA* gene copy number, likely decreasing the metabolic burden associated with high xylose isomerase expression. A synergistic effect was observed in the *isu1*Δ*/zwf1*Δ double mutant, further boosting xylose consumption rates. A diploid derivative (AceY-2n) demonstrated high productivity and robustness in fermentations using hydrolysates from various lignocellulosic feedstocks, highlighting the strain’s potential for industrial-scale applications. These findings reveal novel metabolic targets for strain optimization and offer valuable insights for the rational engineering of yeast platforms for sustainable biofuel and bioproduct production.

## INTRODUCTION

Rising global energy demands and the adverse environmental impacts of fossil fuel consumption have accelerated the development of sustainable technologies. Among these, lignocellulosic biomass has been widely investigated for producing various bioproducts (Antar et al., 2021). Second-generation (2G) technologies, derived from lignocellulosic materials, stand out as a promising sustainable alternative and have been expanding globally, with production facilities now established worldwide (Dos Santos et al., 2016a). Key advancements have been made with the development of robust engineered xylose-fermenting *Saccharomyces cerevisiae* strains, capable of converting xylose into ethanol on an industrial scale. However, major bottlenecks still persist, limiting the full industrial potential of these technologies.

Recent advances in synthetic biology are transforming this field, creating technologies that support the development of viable, large-scale industrial biorefineries. AI-driven genomic metabolic models are being used to optimize pathway design and guide protein-directed evolution, creating highly efficient catalysts that enhance bioprocessing (Gong et al., 2024; Jang et al., 2022). Meanwhile, strategies for modular assembly of multi-gene pathways, coupled with large combinatorial expression libraries to fine-tune metabolic components, enable the streamlined expression of complex biochemical processes (Young et al., 2021). Rational design of complex genetic circuits contributes to alleviating metabolic burden, thereby optimizing metabolic flux in engineered microbes (Xu et al., 2024). Together, these advances provide a robust foundation for scalable bio-based production, enabling biorefineries to meet industrial demands efficiently.

In baker’s yeast, xylose can be converted to xylulose through the heterologous expression of xylose isomerase (*xylA*) (Kuyper et al., 2003; Madhavan et al., 2009). Xylulose is subsequently phosphorylated to xylulose-5-phosphate, entering the non-oxidative phase of the pentose phosphate pathway (PPP). The xylose isomerase pathway is considered more efficient than the two-step XR-XDH pathway in eukaryotes, as the latter pathway often results in cofactor imbalance, leading to the accumulation of fermentation byproducts and compromising yield and productivity (Karhumaa et al., 2007). Engineering *S. cerevisiae* strains with the *xylA* pathway has yielded efficient xylose-converting strains, although optimal performance in xylose fermentation has consistently been achieved only after adaptive laboratory evolution (ALE) (Demeke et al., 2013; Diao et al., 2013; Kuyper et al., 2005; Lee et al., 2014). The evolutionary process has fixed mutations in key components that accelerate xylose fermentation. Interestingly, most mutations described to date in the literature are not obviously related to carbohydrate metabolism, notably including components of iron-sulfur cluster biogenesis (*ISU1* and *CCC1*), components of MAP kinase (MAPK) signaling (*HOG1*, *SSK2*), RAS signaling pathway (*IRA2*), a Golgi Ca^2+^/Mn^2+^ ATPase (*PMR1*), and a heavy metal ion homeostasis protein (*BSD2*) (Dos Santos et al., 2016a; Palermo et al., 2021; Sato et al., 2016; Verhoeven et al., 2017). Understanding the genetic and molecular mechanisms that govern the complex metabolic network driving C5 fermentation is essential. Furthermore, elucidating how these mechanisms relate to the sensing, signaling, and utilization of alternative carbon sources in engineered *S. cerevisiae* is vital for developing microbial platforms suitable for biorefinery applications.

In addition to efficient xylose utilization, high tolerance to fermentation inhibitors is critical for 2G bioproducts. During biomass pretreatment and hydrolysis, compounds are generated that deplete essential cellular energy reserves (e.g., ATP, NADH, NADPH), directly affecting the yield and productivity rates of the strains used (Piotrowski et al., 2014). These inhibitors fall into three main categories: (I) furans (e.g., furfural and HMF); (II) weak acids (e.g., acetic, formic, and levulinic acids); and (III) phenolic compounds (Almeida et al., 2007; Ling et al., 2014). Although solvent-based methods have been investigated for inhibitor removal, these processes increase both production time and cost, limiting economic viability (Jönsson and Martín, 2016; Nascimento et al., 2023; Roque et al., 2019).

Among these inhibitors, acetic acid, derived from the deacetylation of hemicellulose, presents a major challenge, particularly under the low pH conditions typical of fermentation processes. At a pH below its pKa, acetic acid remains undissociated, allowing it to diffuse across cell membranes (Palmqvist and Hahn-Hägerdal, 2000). Once inside the cell, acetic acid dissociates, releasing protons (HlJ), which cannot readily cross the membrane, leading to cytosolic acidification (Mira et al., 2010). Cells counteract this pH drop via membrane ATPase activity (Pma1p), which exports protons at the expense of ATP, directly impairing cell growth and productivity (Almeida et al., 2007; Ullah et al., 2012; Verduyn et al., 1992). Beyond its impact on intracellular pH and cellular energy levels, acetic acid also inhibits nucleic acid synthesis and repair, enzyme activity, macromolecule biosynthesis, and disrupts cell membrane integrity, affecting the proton gradient (Bellissimi et al., 2009; Ding et al., 2013; Fernandes et al., 2005; Piotrowski et al., 2014; Stratford and Anslow, 1998; Ullah et al., 2012).

In this work, we describe how mutations in cell cycle regulation, along with synergistic epistasis in components of the PPP and iron-sulfur metabolism, drive enhanced xylose fermentation and tolerance in industrial *S. cerevisiae*. Our results highlight novel genetic targets for improving microbial strain design, enabling more efficient platforms to meet the demands of industrial-scale biorefineries.

## RESULTS AND DISCUSSION

### Acetic acid accelerates a two-stage evolution for C5 fermentation and tolerance

The haploid strain of *S. cerevisiae* LVY34.4 (Dos Santos et al., 2016b) was constructed using the industrial yeast background of PE-2, which is widely employed in industrial processes due to its high robustness against various fermentation stresses (Basso et al., 2008). This strain was engineered to integrate the *xylA* gene from *Orpinomyces sp.* (Madhavan et al., 2009), *GRE3* deletion, and overexpression of genes from the non-oxidative phase of the pentose phosphate pathway (*RKI1, RPE1, TKL1, TAL1*) and the xylulokinase *XKS1* (Sup. Table S1). LVY34.4 acquired rapid xylose assimilation and fermentation capabilities after an adaptive evolution process that fixed a *Leu132Phe* loss-of-function mutation in *ISU1*, a scaffold protein responsible for de novo iron–sulfur cluster assembly, and introduced a tandem amplification of *xylA*, increasing its copy number and activity. The evolved strain was subsequently selected for a second stage of evolution targeting acetic acid tolerance, a major inhibitor found in lignocellulosic hydrolysates.

The evolution of LVY34.4 for acetic acid tolerance was carried out in sealed bottles under semi-anaerobic conditions, using xylose as the sole carbon source. Acetic acid was initially added at a final concentration of 4 g/L, without pH adjustment, to induce the toxic cellular effects of the undissociated form of the weak acid at low pH (Mira et al., 2010). Successive batch cultures were conducted under the imposed selective conditions until a shortened lag phase and an increased cell growth rate were observed. The acetic acid concentration was then increased to 5 g/L, and the same selection criteria were applied in a new round of experiments, gradually raising the acetic acid concentration to a final level of 8 g/L. As the concentration of acetic acid used increased, there was a decrease in pH, eventually reaching a value below 4 at the highest concentration (Figure 1A).

**Figure 1.**
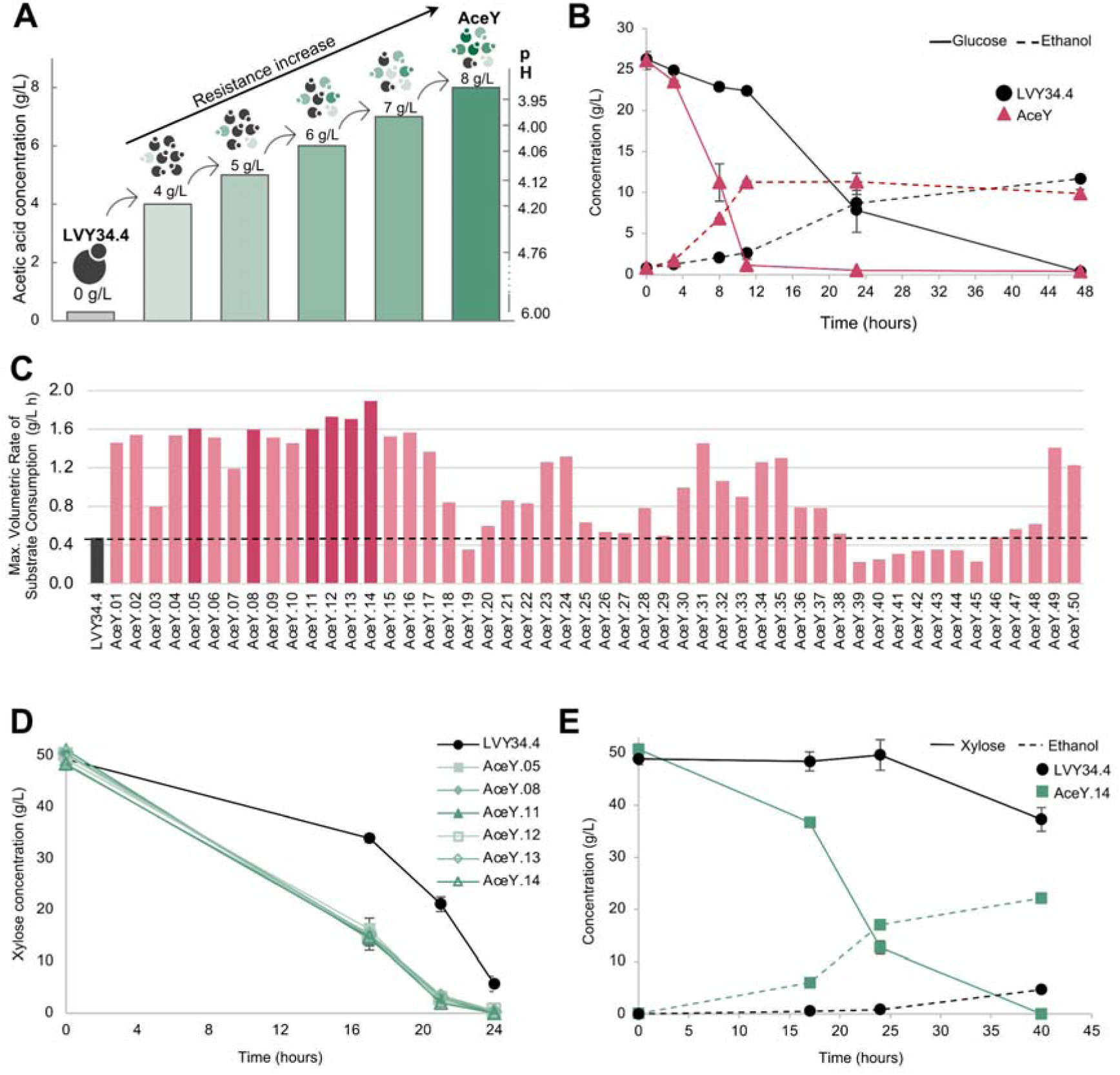
Selection of a strain with enhanced acetic acid tolerance and accelerated xylose fermentation through a second-stage evolution. (A) Graphical representation of the selective pressure conditions, illustrating the acetic acid concentrations added and the corresponding approximate pH values of the medium at each level. (B) Fermentative comparison between the parental strain LVY34.4 and the evolved population AceY in YPD supplemented with 7 g/L acetic acid. (C) Maximum volumetric xylose consumption rate of 50 isolated colonies from the evolved pool in YP medium containing 50 g/L xylose and 5 g/L acetic acid, compared to the parental LVY34.4 (black dashed line). Fermentation was carried out for 31 hours. The top isolates are highlighted in dark pink. (D) Xylose consumption of the top-performing isolated strains compared to the parental LVY34.4 in YP medium with 50 g/L xylose. (E) Fermentative performance of the evolved isolated AceY.14 and its parental strain LVY34.4 in YP medium containing 50 g/L xylose and 4 g/L of acetic acid. All error bars represent the standard deviation of two technical replicates.

The impact of pH on acetic acid toxicity during xylose consumption is well-documented; at low pH, pentose consumption is nearly extinguished in the presence of acetic acid (Bellissimi et al., 2009; Sànchez I Nogué et al., 2013). When the pH drops below the weak acid’s pKa of 4.76, the undissociated form of acetic acid prevails, allowing it to diffuse across the yeast plasma membrane and reduce the cytosolic pH (Palmqvist and Hahn-Hägerdal, 2000). To restore intracellular pH, the cell activates the plasma membrane ATPase Pma1p to expel protons, consuming ATP in the process (Mira et al., 2010). Given that xylose fermentation produces less ATP than glucose fermentation (1.67 vs. 2.0 mol ATP/mol xylose) (McMillan, 1993), the conditions of this evolution strategy are particularly challenging for the cells.

Tolerant mutants were isolated and screened after evolution in media containing 6, 7, and 8 g/L of acetic acid. Cell populations evolved in 6 and 7 g/L acetic acid initially exhibited shorter lag phases during fermentation in YPD supplemented with acetic acid, and resistant clones were isolated. However, after serial transfers in acetic acid-free media, these clones lost their acquired resistance, showing growth patterns similar to the parental strain LVY34.4 (data not shown). Similar results have been described in the literature, where short periods of acetic acid exposure triggered cellular adaptation, increasing resistance to this weak acid (Sànchez I Nogué et al., 2013; Wright et al., 2011); however, without continuous exposure to the inhibitor, this resistance fades (Wright et al., 2011). This transient resistance was likely induced by an epigenetic effect related to histone acetylation, altering global expression patterns to improve fitness under acetic stress, as acetate can act as an epigenetic regulator to promote cell survival (Gao et al., 2016; Salas-Navarrete et al., 2021). A stable resistance phenotype was observed only in mutants isolated from populations evolved in 8 g/L acetic acid. This evolved cell population, termed AceY, showed enhanced sugar consumption and higher ethanol production rates compared to LVY34.4 when grown in YPD supplemented with acetic acid (Figure 1B).

Cells from the AceY population were isolated on solid YNBX medium supplemented with acetic acid. Fifty of the largest colonies were selected and evaluated against the parental strain LVY34.4 in YP medium containing 50 g/L of xylose and 5 g/L of acetic acid. Some isolated clones exhibited a maximum volumetric xylose consumption rate up to three times higher than that of the parental strain (Figure 1C). The evolved strains AceY.05, AceY.08, AceY.11, AceY.12, AceY.13, and AceY.14 showed superior overall performance, prompting a new comparative fermentation among the top isolates (Figure 1D). In the absence of the inhibitor, all isolates demonstrated similar xylose consumption rates, with no significant differences observed. Notably, the selected strains also assimilated xylose more rapidly than LVY34.4 under inhibitor-free conditions (Figure 1D). This finding indicates that adaptive evolution not only enhanced tolerance to inhibitors but also improved xylose consumption rates in the evolved strains.

In the second stage of evolution, acetic acid served as a critical selective agent, amplifying evolutionary pressure and imposing a metabolic constraint that accelerated the emergence of strains with superior xylose consumption capabilities. As a control, a parallel experiment was conducted in triplicate without the addition of acetic acid, and no significant improvement in xylose consumption rate was observed (data not shown). Among the selected strains, AceY.14 demonstrated the highest xylose consumption in the presence of acetic acid, confirming its inhibitor tolerance. Compared to the parental strain LVY34.4, AceY.14 showed enhanced acetic acid tolerance and exhibited faster xylose consumption and ethanol production (Figure 1E). It is worth noting, however, that the resistance level of AceY.14 to acetic acid varied depending on the exposure pattern. When the strain was subjected to continuous acetic acid exposure, it rapidly developed increased resistance. This adaptive response was not observed in the parental strain LVY34.4, suggesting that one or more mutations acquired during evolution enabled AceY.14 to dynamically adjust to acetic acid stress, enhancing tolerance to this weak acid. Therefore, this strain was chosen for further characterization.

### Whole-genome sequencing reveals a fine-tuned adjustment in *xylA* copy number and sparse SNP occurrence

Genomic DNA sequencing of AceY.14 was performed using the Illumina platform, generating 14.4 million paired-end reads of 100 bp each, with a coverage of 120x. The genome was assembled using the SPAdes toolkit (Bankevich et al., 2012), resulting in 214 contigs larger than 1,000 bp, totalling 11.6 Mb of assembled genome and an N50 of 109.6 Kbp. Sequencing reads were aligned to the parental strain LVY34.4, with 88.6% successfully mapping to its genome. Previous molecular and structural analyses of strain LVY34.4 revealed that unequal crossovers involving sister chromatids led to amplifications of the insert containing the *xylA* gene *in tandem* during adaptive evolution on xylose, ultimately resulting in 36 gene copies (Dos Santos et al., 2016b). Despite the improved xylose consumption rate, the 6,237 bp region containing the *xylA* and *LEU2* genes, previously amplified in LVY34.4, did not experience further amplification in AceY.14. Surprisingly, it was reduced by about 20 copies, resulting in about 16 copies of *xylA* integrated near CEN5 in this strain (Figure 2A). As the *xylA* genes are arranged *in tandem*, a recombination event likely occurred at this locus, leading to gene excision and copy number reduction. We hypothesize that this decrease in copy number may help alleviate the metabolic burden associated with high-level expression of this heterologous protein, as overexpression can disrupt normal cellular functions (Glick, 1995; Mao et al., 2024). Several strategies to reduce this metabolic load have emerged, aided by artificial intelligence, genome-scale metabolic models (GEMs), and synthetic biology (reviewed by Mao et al., 2024). A recent ALE experiment in xylose also reported a decrease in *xylA* copy number after a prolonged period, although it stabilized with a higher copy number than in our strain, decreasing from approximately 134 copies to 64 copies (Zhang et al., 2024). In a newly conducted long-term evolution experiment using the industrial PE-2 strain to construct improved xylose-fermenting strains, a ∼16-fold amplification of *xylA* was also observed across 10 different populations (personal communication), suggesting that this may represent an optimal copy number for balancing efficient xylose fermentation with cellular metabolic demands in PE-2 background strains.

**Figure 2.**
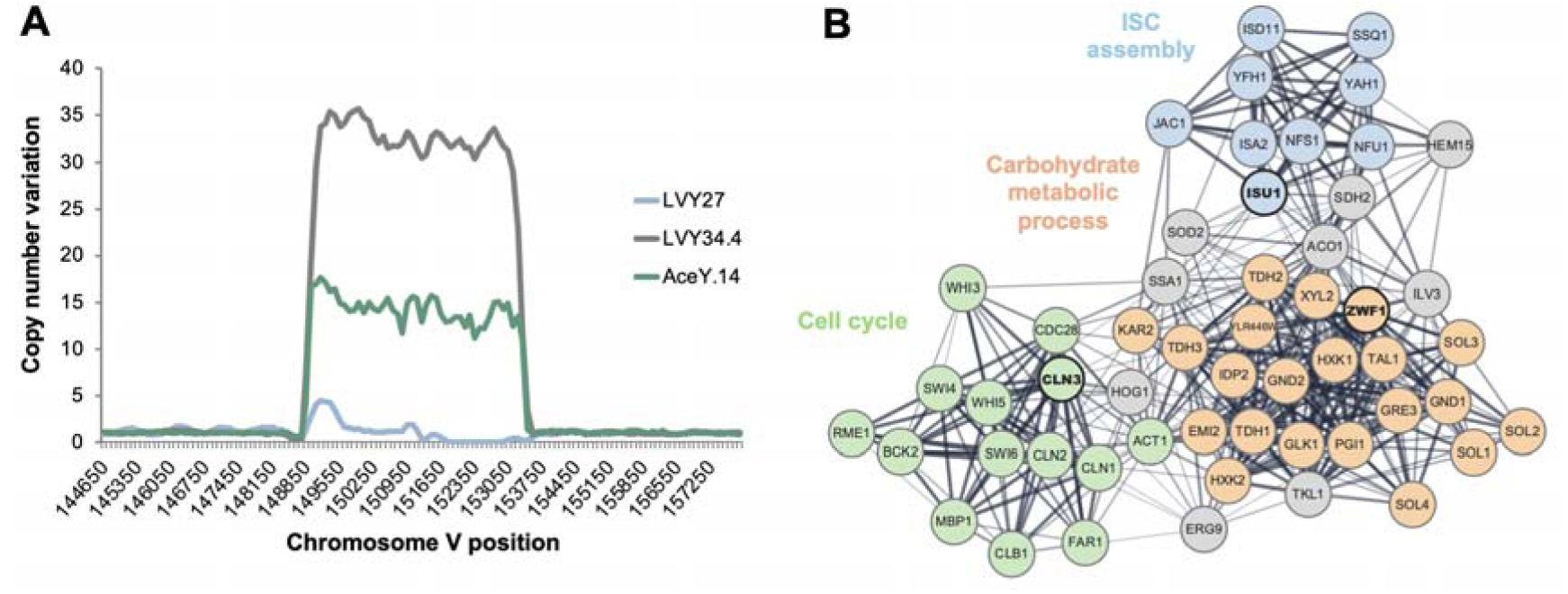
*xylA* copy number variation and gene interaction network of mutants. **(A)** Normalized coverage of chromosome V showing the copy number variation of *xylA* in the engineered strain LVY27 (blue), the evolved parental strain LVY34.4 used in this study (gray), and the acetic acid evolved strain AceY.14 isolated in this study (green). LVY27, which contains only one copy of *xylA*, underwent ALE on xylose, resulting in the strain LVY34.4 (Dos Santos et al., 2016a), carrying 36 copies of this gene. The AceY.14 strain exhibited a reduction of approximately 20 copies of *xylA* compared to its parental strain LVY34.4. **(B)** Protein-protein interaction network of the novel mutated genes *CLN3* and *ZWF1* found in the AceY.14, as well as the *ISU1* mutation already present in the parental strain LVY34.4. The network was constructed using Cytoscape 3.10.3 (Shannon et al., 2003) and data collected from the stringApp (v2.1.1), incorporating 50 additional interactions with a confidence score threshold of 0.4. Genes were grouped according to Gene Ontology (GO) enrichment: blue - Iron-Sulfur Cluster (ISC) assembly (GO:0016226); orange - Carbohydrate Metabolic Process (GO:0005975); green - Cell Cycle (GO:007049).

The identification of single nucleotide polymorphisms (SNPs) revealed a small number of mutations, despite the extended duration and strong selective pressure of the second stage of evolution (Table 1). Nucleotide changes identified in the *YRF1*, *NUD1,* and *HXT6* genes resulted in synonymous substitutions. Interestingly, the modification of five bases in the hexose transporter *HXT6*, spaced within 14 nucleotides, did not change the amino acid sequence of the transporter. A point mutation was also identified in the left arm of the telomeric region of chromosome VIII. The only amino acid substitution occurred in the gene encoding the glucose-6-phosphate dehydrogenase *ZWF1*, which catalyses the first step of the pentose phosphate pathway. This mutation resulted in the replacement of glutamic acid with aspartic acid at position 191. Additionally, a frameshift mutation (T556fs) was identified in the *CLN3* gene, which encodes a cyclin involved in cell cycle regulation, introducing a premature stop codon.

**Table 1.**
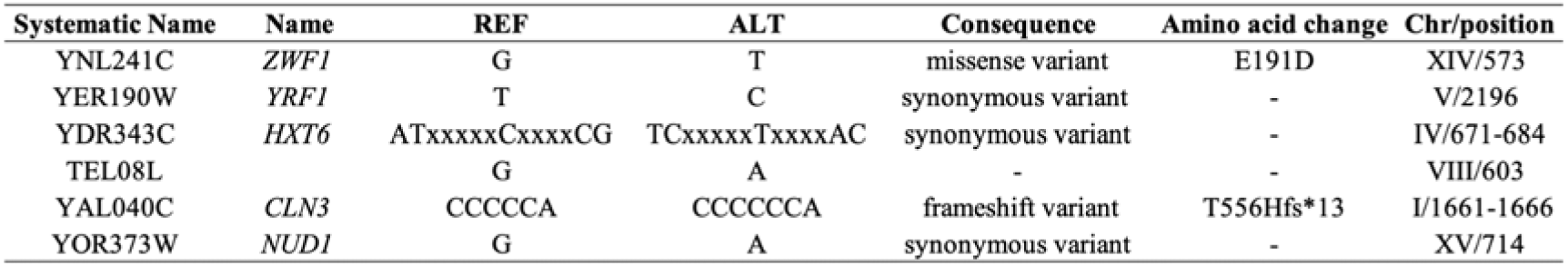
Mutations identified in the AceY.14 strain.

A protein-protein interaction network analysis did not reveal any direct links between these mutated genes or associated metabolic pathways, nor any obvious connection to iron-sulfur metabolism, despite the parental strain LVY34.4 carrying a loss-of-function mutation in *ISU1* (Figure 2B). Aside from *ZWF1*, there are no clear connections between these mutations and xylose fermentation or carbohydrate metabolism, highlighting the adaptive plasticity of yeast metabolic networks in response to evolutionary pressures. Mutations in *ZWF1* and *CLN3* were then further investigated to assess their potential roles in the enhanced xylose assimilation and acetic acid resistance observed in this strain.

### Investigating the role of *ZWF1* mutations on xylose utilization in *xylA*-expressing strains

The strain C5TY (C5 Trial Yeast), also known as LVY59, was used for functional analysis of nucleotide mutations through reverse engineering. This strain is derived from LVY34.4 and was constructed by reverting the *ISU1* point mutation in LVY34.4 back to the wild-type allele (Dos Santos et al., 2016a). As this strain carries the genetic modification required for xylose consumption (overexpression of xylulokinase and PPP genes, and multiple copies of *xylA*) without any additional mutations, it provides an ideal platform for testing the effects of candidate genes on xylose metabolism. The strategy used to evaluate the effect of each mutation was based on obtaining a knockout strain to assess the impact of gene loss-of-function, followed by reversing the gene deletion by integrating the target gene carrying the specific SNP using a CRISPR system. Deletion mutants derived from the C5TY strain were constructed by gene disruption using the *MX* deletion cassettes (Goldstein and McCusker, 1999; Wach et al., 1994), and the specific point mutations were integrated by replacing the deletion cassette to the mutant gene using the CRISPR/Cas9 EasyGuide toolkit (Jacobus et al., 2022) and gRNAs targeting the *MX* fragment. This approach enabled precise and targeted modification of individual nucleotide sequences, providing a powerful method for dissecting the influence of specific mutations on metabolic performance.

The *ZWF1* gene encodes the yeast glucose-6-phosphate (G6P) dehydrogenase enzyme (EC 1.1.1.49), which catalyses the first step of the oxidative phase of the pentose phosphate pathway (PPP), a key metabolic route responsible for regenerating NADPH from NADPL through an oxidation-reduction reaction (Figure 3A). During this process, one carbon is lost as COL for every two NADPH molecules regenerated. This step is crucial for NADPH production, an essential molecule for various cellular functions and for maintaining redox balance (Celton et al., 2012; Dos Santos et al., 2025). Interestingly, mutants with a deletion of *ZWF1* demonstrated surprisingly better performance in xylose fermentation compared to the C5TY control strain. Strains harbouring the point mutation *zwf1^E191D^* also showed improved performance, though not to the same extent as the knockout strain (Figure 3B). Specifically, the *zwf1*Δ and *zwf1^E191D^* strains exhibited increases in maximum productivity rates of approximately 97% and 66%, respectively, compared to the C5TY control (Table 2). Furthermore, strains lacking *ZWF1* also showed the lowest xylitol yield (Table 2 and Figure 3C). As xylitol is a known inhibitor of xylose isomerase activity (Yamanaka, 1969), this result is consistent with previous findings that *ZWF1* deletion decreases xylitol and increases ethanol yields (Jeppsson et al., 2002; Verho et al., 2003). These studies also reported that acetate levels rise in response to the redox cofactor imbalance caused by *ZWF1* loss of function, which was similarly observed in *zwf1*Δ strains in the present study (Table 2 and Figure 3C). Although the strain carrying the point mutation exhibited a decrease in xylitol yield, it did not show a significant increase in acetate production (Table 2).

**Figure 3.**
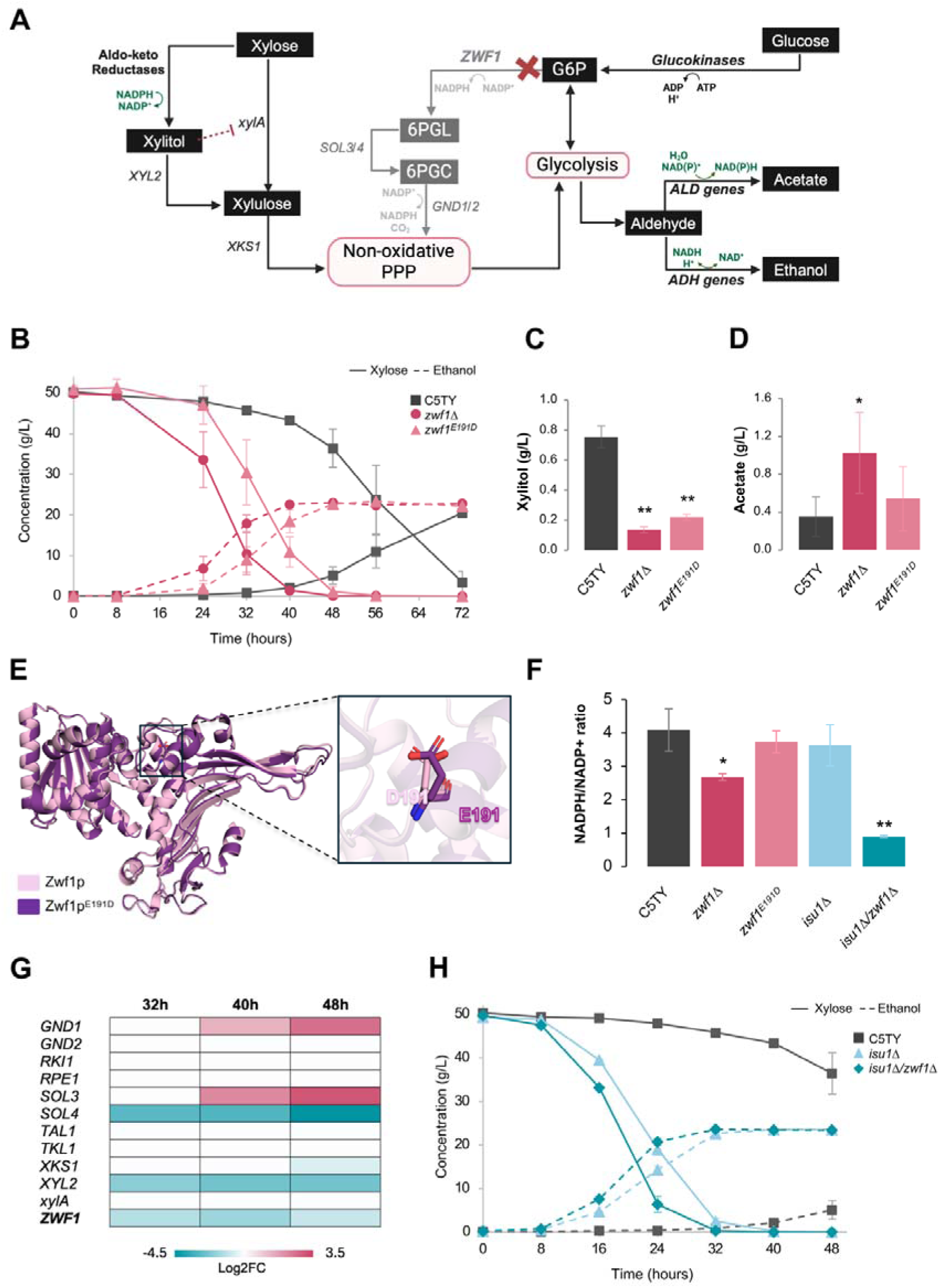
Assessing the role of *ZWF1* mutations on xylose metabolism in yeast. (A) Simplified overview of *ZWF1’*s role in the oxidative PPP and the effects of its deletion on xylose fermentation. *ZWF1* encodes the enzyme catalyzing the first step of the oxidative phase of the pentose phosphate pathway (PPP) and its deletion reduces NADPH regeneration via this pathway (indicated as faded). This NADPH depletion decreases the activity of NADPH-dependent endogenous aldo-keto reductases, reducing the conversion of xylose to xylitol, a known inhibitor of xylose isomerase (*xylA*). Consequently, acetate production increases to provide an alternative source of NADPH. (B) Fermentative performance, (C) xylitol production, and (D) acetate production of mutants with *ZWF1* deletion or point mutation, compared to the control strain C5TY, grown in rich YP medium with 50 g/L xylose. Xylitol and acetate production are shown at 72 hours of fermentation (final time point). Data represent the mean ± standard deviation of two independent biological replicates, each performed with technical triplicates. (E) Structural superimposition of *Sc*Zwf1 predicted model showing that Glu191Asp replacement likely disrupts interactions required for activity. (F) NADPH/NADP^+^ ratio of *ZWF1* mutants and the control on xylose, represented by mean ± standard deviation of technical triplicates. (G) Time-course profile of pentose phosphate pathway genes of C5TY during fermentation in YP with 50 g/L of xylose. Color degrees show up- (pink red) and downregulation (green blue) of genes to average Log2 fold change values in each time-point compared with the sample of 8 h. Data from YeastC5 platform (Palermo et al., 2021) (H) Fermentative performance of the double deletant *isu1*Δ*/zwf1*Δ *in* rich media YP with 50 g/L of xylose compared to the control and the single mutant *isu1*Δ. Data represent the mean ± standard deviation of technical replicates. * p < 0.05, ** p < 0.01

**Table 2.**
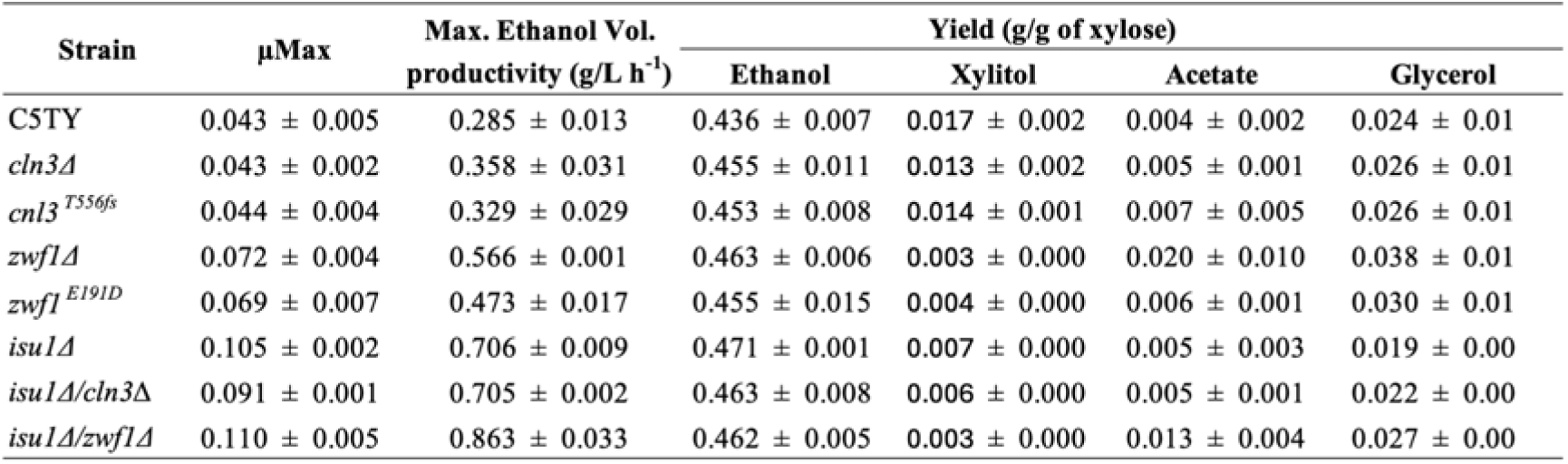
Fermentation performance of mutant strains and the control C5TY in YP medium with 50 g/L of xylose.

| Strain | $\mu_{\text{Max}}$ | Max. Ethanol Vol.<br>productivity (g/L h<sup>-1</sup>) | Yield (g/g of xylose) | | | |
| --- | --- | --- | --- | --- | --- | --- |
|  |  |  | Ethanol | Xylitol | Acetate | Glycerol |
| C5TY | 0.043 ± 0.005 | 0.285 ± 0.013 | 0.436 ± 0.007 | 0.017 ± 0.002 | 0.004 ± 0.002 | 0.024 ± 0.01 |
| *cln3Δ* | 0.043 ± 0.002 | 0.358 ± 0.031 | 0.455 ± 0.011 | 0.013 ± 0.002 | 0.005 ± 0.001 | 0.026 ± 0.01 |
| *cln3<sup>T556fs</sup>* | 0.044 ± 0.004 | 0.329 ± 0.029 | 0.453 ± 0.008 | 0.014 ± 0.001 | 0.007 ± 0.005 | 0.026 ± 0.01 |
| *zwf1Δ* | 0.072 ± 0.004 | 0.566 ± 0.001 | 0.463 ± 0.006 | 0.003 ± 0.000 | 0.020 ± 0.010 | 0.038 ± 0.01 |
| *zwf1<sup>E191D</sup>* | 0.069 ± 0.007 | 0.473 ± 0.017 | 0.455 ± 0.015 | 0.004 ± 0.000 | 0.006 ± 0.001 | 0.030 ± 0.01 |
| *isu1Δ* | 0.105 ± 0.002 | 0.706 ± 0.009 | 0.471 ± 0.001 | 0.007 ± 0.000 | 0.005 ± 0.003 | 0.019 ± 0.00 |
| *isu1Δ/cln3Δ* | 0.091 ± 0.001 | 0.705 ± 0.002 | 0.463 ± 0.008 | 0.006 ± 0.000 | 0.005 ± 0.001 | 0.022 ± 0.00 |
| *isu1Δ/zwf1Δ* | 0.110 ± 0.005 | 0.863 ± 0.033 | 0.462 ± 0.005 | 0.003 ± 0.000 | 0.013 ± 0.004 | 0.027 ± 0.00 |

The Zwf1p protein of PE-2 lacks a glutamic acid at position 59 compared to the reference genome strain S288C, similar to some alternative reference strains which also lack this amino acid (e.g., Sigma1278b, SK1, Y55). On the other hand, the point mutation E191D occurs in a conserved nine-residue peptide (RIDHYLGKE) (Fig. 3E, Fig. S1), which has been described as containing key residues in the G6P binding site of the human homolog protein (Kotaka et al., 2005). Based on our results, the substitution in the mutant strain of glutamic acid by aspartic acid—both amino acids with negatively charged side chains—likely decreased Zwf1p activity rather than eliminating its function, as the deletion and point mutation exhibited different phenotypes under the conditions tested. Indeed, the substitution of glutamate by a residue with a shorter lateral chain, such aspartate, can disrupt the interactions required for substrate positioning and catalysis, lowering the activity of Zwf1p (Fig. 3E).

Mutations in the *ZWF1* gene have been explored in strains carrying the oxidoreductive (XR/XDH) pathway to address cofactor imbalance, with strategies such as *ZWF1* disruption combined with other genetic modifications to increase the NADP^+^ pool for the XDH enzyme (Jeppsson et al., 2002; M Jeppsson et al., 2003; Verho et al., 2003). In contrast, *ZWF1* deletion and other PPP disruptions reduce the supply of NAD(P)H for detoxifying inhibitors like furfural and HMF, thereby increasing sensitivity to these compounds (Gorsich et al., 2006; Marie Jeppsson et al., 2003; Lewis Liu et al., 2009; Dos Santos et al., 2025). To assess the mutation’s effect on acetic acid resistance and the NADPH/NADP^+^ ratio, we tested the strain’s growth in 4 g/L of acetic acid at pH ∼4.3 (below the pKa) (Fig. S2 and S3) and measured the NADPH/NADP^+^ ratios on xylose (Fig. 3F). The *zwf1*Δ and *zwf1^E191D^* mutants showed no significant reduction in growth on YPX plates with acetic acid compared to the control (Fig S2); however, *zwf1*Δ displayed reduced growth in liquid YPD media with acetic acid (Fig. S3A). Furthermore, strains with *zwf1*Δ were the only ones showing a significant decrease in the NADPH/NADP+ ratio compared to the control (Fig. 3F), confirming previous findings reported by Qiu et al. (2024).

The modification of *ZWF1* expression has also been described recently in a strain harbouring the xylose isomerase pathway (Qiu et al., 2024). In contrast to the present study, Qiu et al. (2024) observed impaired growth and sugar consumption following *ZWF1* deletion, which was recovered by the introduction of heterologous NADP^+^-dependent glyceraldehyde- 3-phosphate dehydrogenase genes, highlighting the critical role of NADPH availability in supporting central metabolism. Curiously, the *ZWF1* mutants in this study exhibited enhanced xylose consumption without growth reduction on glucose or xylose (Fig. S3B and S4). Qiu et al. (2024) hypothesized that blocking glucose entry into the PPP increases xylose utilization. Indeed, previous findings from our group have shown that transcription data of xylose isomerase-expressing strains exhibit reduced *ZWF1* expression (Fig. 3G) (Palermo et al., 2021), in agreement with findings from (Qi et al., 2015). Additionally, metabolic flux analysis of non-xylose-fermenting *ZWF1* deletants reported a reversed flux in the non-oxidative PPP, supporting the production of the building blocks erythrose-4-phosphate and ribose-5-phosphate (Blank et al., 2005). These data suggest that the increased xylose consumption observed in the *ZWF1* mutants may be a combined effect of adaptation within the non-oxidative PPP in response to oxidative phase inactivation, and the low xylitol yield, which results in reduced *xylA* inhibition, thereby increasing flux through this pathway and consequently enhancing xylose consumption.

### *ZWF1* loss-of-function has a synergistic effect with *ISU1* metabolism in increasing the xylose consumption rate

The xylose-fermenting strain LVY34.4, used as the background for the second stage of evolution, was previously evolved on xylose, acquiring the *Leu132Phe* mutation in the *ISU1* gene, which encodes a protein involved in mitochondrial iron-sulfur cluster biogenesis (Garland et al., 1999; Lill et al., 2012). This mutation, along with multiple copies of *xylA,* has been reported to enhance xylose fermentation in this strain (Dos Santos et al., 2016b). To investigate whether *ZWF1* loss-of-function could further improve xylose fermentation in the presence of the *ISU1* mutation, we additionally deleted *ISU1* in the *ZWF1* deletant strains and in the control strain C5TY, since *ISU1* deletion leads to a xylose consumption rate similar to that of the point-mutated strain (Dos Santos et al., 2016b).

The *isu1*Δ*/zwf1*Δ double mutant exhibited an enhanced xylose consumption profile, with volumetric productivity increasing by 22% compared to *isu1*Δ (Figure 3H, Table 2). This fermentative performance was superior to that identified by the two single mutants *zwf1*Δ and *isu1*Δ, suggesting a synergistic interaction between the two mutations that positively affects xylose consumption.

The relationship between *ISU1* deletion and increased xylose consumption has been well described in the literature (Sato et al., 2016; Dos Santos et al., 2016b; Osiro et al., 2019; Lee et al., 2021; Palermo et al., 2021), and these studies have also shown the synergistic effect of this mutation with other genes. As a metalloenzyme, xylose isomerase (XI) benefits from the increased cellular iron availability caused by *ISU1* deletion, potentially enhancing XI activity and xylose consumption. Similar effects have been observed with mutations that increase cytosolic metal concentrations (Palermo et al., 2021). The observed epistatic interaction in the *isu1*Δ*/zwf1*Δ double mutant likely reflects a combination of improved XI activity due to *ISU1* deletion and reduced xylitol yield, along with expanded flux through the non-oxidative PPP, driven by metabolic remodelling in response to oxidative phase inactivation caused by *ZWF1* deletion.

### Impact of the G1 cyclin *CLN3* on cell cycle progression, xylose fermentation, and acetic acid resistance

The second point mutation present in AceY.14 is in the *CLN3* gene, which encodes a G1 cyclin responsible for initiating the cell cycle. In the early G1 phase, Cln3p forms a complex with the cyclin-dependent kinase Cdc28p and acts as an upstream activator of other G1 cyclins, Cln1p and Cln2p, through the derepression and activation of the transcription factors SBF and MBF, respectively, thereby promoting cell cycle progression (Fig. 4A) (Costanzo et al., 2004; De Bruin et al., 2004). Loss of *CLN3* causes delayed cell cycle progression and increased cell size due to elongation of the G1/S transition (Talia et al., 2007; Teufel et al., 2019; Zhang et al., 2002). Deletion of *CLN3* led to an increase in cell size in the mutant strains (Fig. 4B), a trait also observed in the evolved strain AceY.14 (Fig. S5).

**Figure 4.**
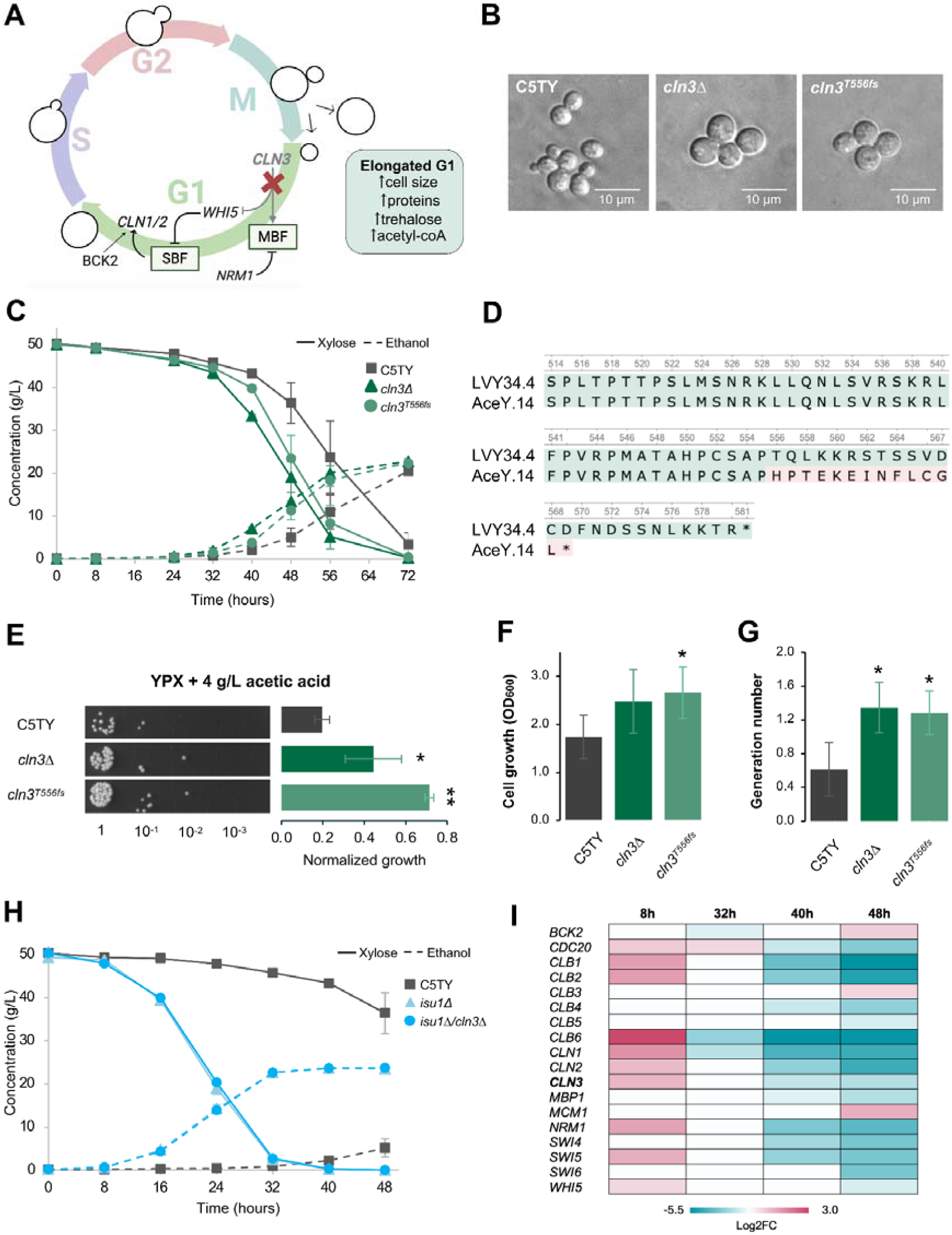
*CLN3* mutations affect xylose fermentation and resistance to acetic acid. (A) Simplified overview of the *CLN3* role in cell cycle progression. *CLN3* encodes a cyclin responsible for initiating the cell cycle, and its deletion causes elongation of the G1/S transition, leading to increased cell size and enhanced production of cellular metabolites, proteins, and storage carbohydrates. (B) Differential interference contrast (DIC) microscopy of the control and mutant strains of *CLN3*. (C) Fermentative performance of mutants with either deletion or point mutation of *CLN3* compared to the control C5TY in rich media YP with 50 g/L of xylose. Data represent the mean ± standard deviation of two independent biological replicates, each performed in technical triplicates. (D) Sequence alignment of amino acids between LVY34.4 and AceY.14, starting from position 514. The beginning of the frameshift is highlighted in red, and an asterisk denotes the stop codon. (E) Growth on solid YP plates containing 20 g/L xylose and 4 g/L acetic acid. The graph shows the mean ± standard deviation (n = 3) of cellular growth, normalized to growth on plates without acetic acid, using cells adjusted to an OD of 1. Growth was quantified using ImageJ software. (F) Cell growth (OD_600_) and (G) generation number of mutant strains compared to the control after 6 h of cultivation in liquid YP media with 20 g/L of glucose, supplemented with 4 g/L of acetic acid, without pH adjustment (pH ∼4.3). Data represent mean ± standard deviation of two independent biological replicates, each performed in technical duplicates. (H) Fermentative performance of the double deletant *isu1*Δ*/cln3*Δ in rich media YP with 50 g/L of xylose compared to the control and the single mutant *isu1*Δ. Data represent the mean ± standard deviation of technical triplicates. (I) Time-course profile of cell cycle genes of *isu1*Δ compared to the respective time-point of C5TY during fermentation in YP with 50 g/L of xylose. Color degrees show up- (pink red) and downregulation (green blue) of genes to average Log2 fold change values in each time-point compared with the sample of 8 h. Transcriptome data were extracted from the YeastC5 platform (Palermo et. al, 2021). * p < 0.05, ** p < 0.01

Although *CLN3* mutants (both the knockout and point mutation) exhibited a growth pattern similar to the parental strain (Fig. S4), their xylose consumption and ethanol production rates were higher than the control (Fig. 4C, Table 2), with no significant differences between the deletion and the frameshift mutant *cln3^T556fs^*. The frameshift at residue T556 of Cln3p identified in this study generates a premature stop codon, resulting in a protein that lacks the last 12 amino acids of the wild-type version (Fig. 4D). Cln3p is a highly unstable protein, and its C-terminal region has been reported to regulate the protein stability, as truncated proteins are more stable than their full-length counterparts and exhibit a small-cell phenotype due to accelerated cell cycle progression (Cross and Blake, 1993; Tyers et al., 1992; Yaglom et al., 1995). However, truncation of the last 20 amino acids has been associated with a loss of protein function, as it results in the large-cell size phenotype similar to *CLN3* deletion (Yaglom et al., 1995). Our results suggest that the *cln3^T556fs^* mutant leads to a loss of function, as it results in the large-cell phenotype observed in the *CLN3* deletion mutant (Fig. 4B), and displays the same increase in xylose consumption rate also present in the *cln3*Δ mutant (Fig. 4A).

In addition to altered cell size, elongation of G1 phase has been associated with trehalose accumulation (Paalman et al., 2003). Trehalose is a storage carbohydrate well known for its protective role against different stresses, such as high temperature, freeze-thaw cycles, high ethanol concentrations, desiccation, and osmotic and oxidative stress (Benaroudj et al., 2001; Chen and Gibney, 2022; Eleutherio et al., 2015; Mahmud et al., 2010). Indeed, our *CLN3* mutants showed slightly increased tolerance to acetic acid compared to the control in agar plates and liquid media (Fig. 4E-G and Fig. S3), suggesting that the T556fs variant in *CLN3* was primarily responsible for the enhanced growth observed in the AceY.14 strain under acetic acid stress. Genes promoting trehalose accumulation have previously been shown to increase resistance to acetic acid (Yoshiyama et al., 2015). Interestingly, a recent ALE experiment using high ethanol concentrations identified evolved strains with enriched mutations in genes related to trehalose metabolism and the cAMP/PKA pathways (Jacobus et al., 2024). Additionally, *CLN3* deletion has been reported to increase acetyl-CoA production (Hao et al., 2024), which could trigger epigenetic effects via histone acetylation, enhancing stress adaptation (Gao et al., 2016; Salas-Navarrete et al., 2021). Together, these findings indicate that trehalose and/or acetyl-CoA accumulation may serve as adaptive mechanisms in *CLN3* mutants exposed to stressful environmental conditions. Therefore, engineering genes involved in these metabolic pathways could improve yeast tolerance to acetic acid.

Similarly to our exploration of *ZWF1* mutations, the potential synergistic effect with the iron-sulfur component *ISU1* was also investigated. The double mutant *cln3*Δ*/isu1*Δ showed no change in the xylose fermentation profile compared to the single mutant *isu1*Δ (Fig. 4H, Table 2), suggesting that *CLN3* deletion does not further enhance xylose metabolism when combined with the *ISU1* mutation. In the *isu1*Δ mutant, genes associated with the cell cycle display a dynamic expression profile, marked by upregulation during active fermentation and subsequent downregulation after xylose is depleted (Fig. 4I), reflecting their involvement in the control of cell division.

Although the role of the *CLN3* gene in improving the xylose consumption rate remains unclear, a previous study also found a link between cell cycle arrest and enhanced xylose utilization. Wei et al. (2018) observed that *NRM1* overexpression increased xylose consumption in *S. cerevisiae* strains, leading to a 35.4% increase in the specific xylose consumption rate in mixed media (20 g/L xylose and 20 g/L glucose). *NRM1* has an opposite role to *CLN3*, as it binds to the transcription factor MBF in the late G1 phase and acts as a transcriptional repressor (De Bruin et al., 2006). *NRM1* overexpression also results in increased cell size, likely due to cell cycle arrest (Travesa et al., 2013). The cell cycle is regulated by numerous nutrient and stress signalling pathways (Adler et al., 2022), some of which have been explored in the context of xylose metabolism—particularly genes controlled by the sugar sensing cAMP/PKA and the High Osmolarity Glycerol (HOG) pathways (Dos Santos et al., 2016b; Sato et al., 2016; Myers et al., 2019; Osiro et al., 2019; Wagner et al., 2019, 2023). Altogether, these findings highlight the complexity of the metabolic network governing xylose fermentation and its integration with broader cellular processes.

### Fermentation of lignocellulosic hydrolysates for sustainable biofuel production

Ethanol biorefineries typically employ robust diploid organisms such as PE-2 and CAT-1, or polyploid strains like Ethanol Red®, due to their greater resilience and tolerance to various stressors (Basso et al., 2008; Favaro et al., 2019). In lignocellulosic biorefineries, in addition to withstanding the inhibitors generated during hydrolysate production from lignocellulosic materials, strains must also efficiently utilize multiple sugar types—including xylose—to support the viable production of lignocellulosic-based bioproducts (Yu et al., 2025). Generally, diploid or polyploid strains demonstrate superior resistance to a range of industrial stresses, a crucial trait for optimizing the conversion of biomass-derived sugars into valuable products such as biochemicals, biofuels, and biopolymers. This adaptability makes diploid and polyploid strains ideal candidates for high-stress industrial environments (Sun et al., 2022; Zhang et al., 2017). Based on these requirements, we self-crossed AceY.14 to generate a diploid strain, designated AceY-2n, which was used for the fermentation of hydrolysates derived from four different biomass types: sugar cane straw, sugar cane bagasse, energy cane bagasse, and eucalyptus. These biomass sources were selected to produce distinct hydrolysates with varying concentrations of inhibitors and sugars, thereby challenging the AceY-2n strain under different conditions that mimic real-world industrial settings.

Cellobiose was present at concentrations ranging from 5 to 7 g/L across all hydrolysates, with the glucose produced from its breakdown quantified and included in the total sugar content calculation. The evolved diploid strain AceY-2n was inoculated at a low initial cell concentration of 0.25 g CDW/L to assess the tolerance acquired by the strain. Furfural and HMF were detected in low amounts (Table S2) and were completely consumed by the end of fermentation.

The eucalyptus hydrolysate exhibited the highest acetic acid concentration, reaching 6.48 g/L (Table S2). Despite the elevated acetic acid levels and the presence of other inhibitors in this hydrolysate, glucose was completely consumed within 15 hours, and xylose remained at a residual concentration of 2.5 g/L after 42 hours. In all hydrolysates, xylose consumption began only after glucose was nearly depleted, as expected due to glucose-induced catabolite repression (Gancedo, 1998). High ethanol yields were observed in all fermentations using the four different biomass types, with the highest yield of 0.42 g/g obtained from sugarcane straw (Figure 5, Table S3).

**Figure 5.**
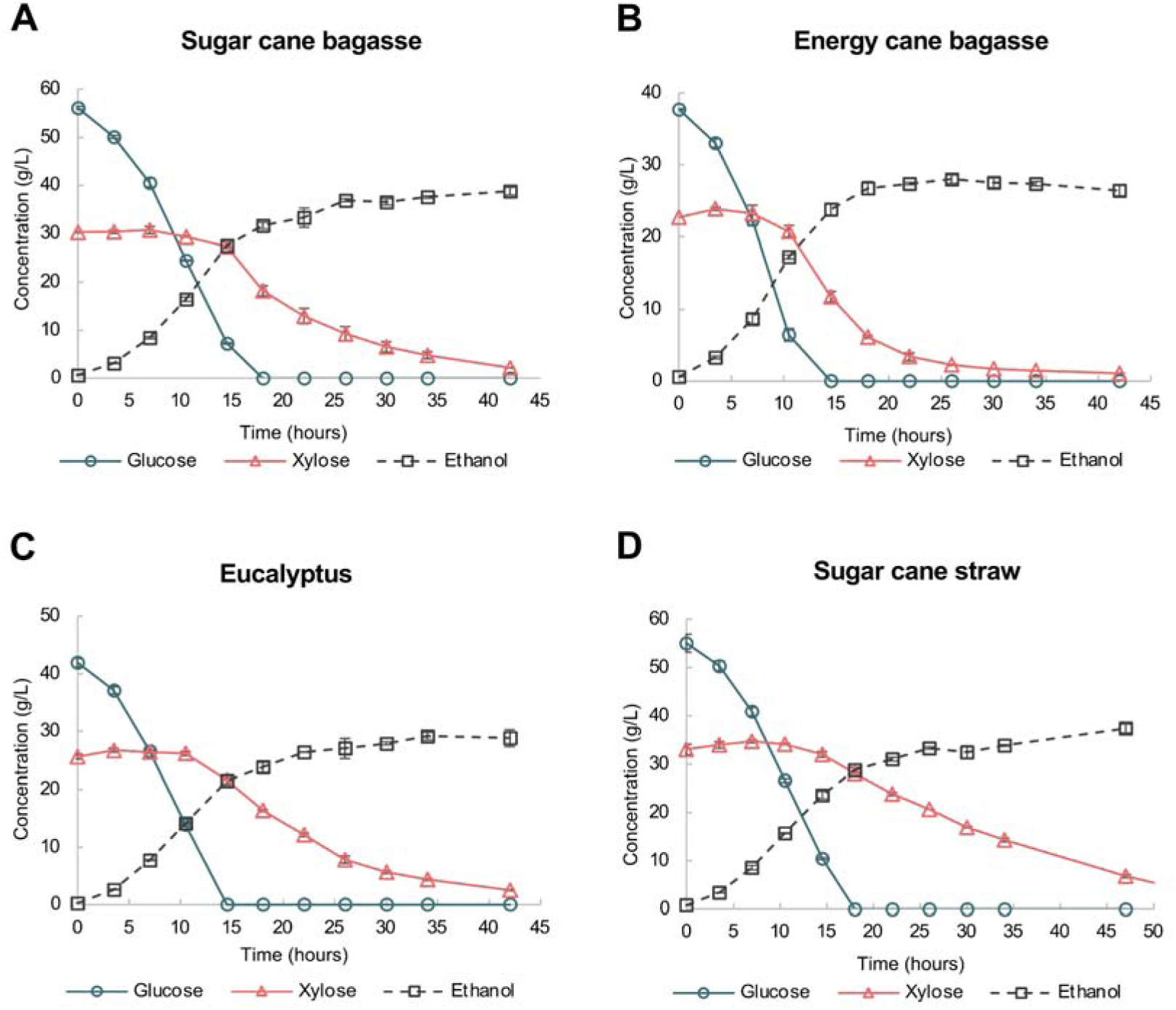
AceY-2n fermentation in different hydrolysates. The fermentative performance of diploid strain AceY-2n was assessed in hydrolysates from sugar cane bagasse (A), energy cane bagasse (B), eucalyptus (C), and sugar cane straw (D). All fermentations were performed in triplicate and the standard deviation is shown by the error bars.

The AceY-2n strain exhibited superior overall performance in both sugarcane and energy cane bagasse hydrolysates, with notably higher volumetric productivity observed in the latter. These results highlight the strain’s ability to efficiently tolerate inhibitors and ferment sugars derived from four distinct biomass sources, underscoring its potential for large-scale biofuel production in industrial settings.

### Conclusions

This study highlights the potential of evolutionary strategies to enhance the robustness of *S. cerevisiae* strains for lignocellulosic-based biorefineries. It also uncovers novel metabolic pathways and genetic determinants that contribute to improved xylose fermentation and inhibitor tolerance in yeast. Through a two-stage evolution process, the evolved strain AceY.14 was isolated, displaying an increased xylose consumption rate and enhanced tolerance to acetic acid. Whole-genome analyses revealed mutations in the cyclin G1 (*CLN3*) and glucose-6-phosphate dehydrogenase (*ZWF1*) genes, with reverse engineering confirming these mutations as key contributors to the observed phenotypic improvements. Specifically, the *cnl3^T556fs^* mutation increased cell size and acetic acid tolerance, potentially through trehalose accumulation or epigenetic adaptation, while the *zwf1^E191D^* mutation boosted xylose consumption, likely by reducing xylitol production and increasing flux through the non-oxidative phase of the pentose phosphate pathway. Notably, the *isu1*Δ*/zwf1*Δ double mutant exhibited an even greater increase in xylose consumption, demonstrating a synergistic epigenetic interaction between the two metabolic pathways. The diploid strain AceY-2n showed high tolerance to inhibitors present in lignocellulosic hydrolysates, efficiently converting sugars to ethanol with high yield and productivity. Overall, this study identifies *CLN3* and *ZWF1* as promising genetic targets for strain improvement and provides new insights into the molecular mechanisms governing xylose metabolism and inhibitor resistance in *S. cerevisiae*. These findings lay the groundwork for the rational engineering of robust yeast strains for the industrial production of second-generation bioproducts.

## METHODS

### Strains and cultivation conditions

The *S. cerevisiae* strains utilized in this study are described in Table S1. The haploid spore LVYA1 is derived from the industrial strain PE-2 (CBMAI 0959). The strains were cultivated in YP medium (10 g/L yeast extract, 20 g/L peptone from gelatin), supplemented with 20 g/L D-glucose (YPD), 50 g/L of xylose (YPX) or 20 g/L of galactose (YPG), and incubated at 30°C, 200 rpm. Colony isolation from the evolved cell pools was performed on YNB medium (6.7 g/L yeast nitrogen base without amino acids), supplemented with 1 g/L drop-out, 20 g/L xylose, 20 g/L agar, and 5 g/L acetic acid. Antibiotics hygromycin (200 mg/L), zeocin (30 µg/mL), geneticin (200 mg/L), and nourseothricin (100 mg/L) were added when necessary for the selection of mutant strains.

Plasmid cloning was performed using *Escherichia coli* DH5α grown in LB medium with 100 mg/L ampicillin.

### Second stage of adaptive evolution

The evolution of the xylose-fermenting strain LVY34.4 was conducted under semi-anaerobic conditions in a sealed 100 mL bottle with a working volume of 80 mL. Evolutions were performed through successive batch cultures in YP medium supplemented with 50 g/L xylose and an initial concentration of 4 g/L acetic acid. No pH adjustment was made during evolution. Each batch was initiated at approximately OD_600_ 2 and transfer to a new medium was performed when it reached approximately OD_600_ 6. The acetic acid concentration was increased after 30 successive transfers, varying from 4 to 8 g/L (Figure 1A). If growth was not observed after inoculation in a new and higher concentration of acetic acid, about 10 to 20 additional transfers were performed in medium with the previous inhibitor concentration. Evolutions were initially conducted in triplicate. However, due to greater growth in one of the experiments, the others were abandoned when the concentration reached 6 g/L. To isolate evolved strains, an aliquot of the evolved pool was inoculated on solid YNB medium containing xylose as the carbon source and supplemented with acetic acid. After incubation at 30°C, colonies showing the largest size were isolated for further characterization.

### Fermentations conditions

Fermentation assays were performed in YPD medium (20 g/L glucose) supplemented with 7 g/L acetic acid to test the evolved cell pool. Exactly 1 mL of the evolved pool was cultivated in YPD overnight to prepare the inoculum for fermentation, starting the culture with an OD_600_ of approximately 1.0. There was no pH control or adjustment, and the temperature and agitation speed were kept constant at 30°C and 200 rpm, respectively, in 100 mL sealed bottles with a working volume of 70 mL.

For isolated colonies and reverse-engineered strains, fermentation was conducted in YP medium supplemented with 50 g/L xylose (YPX), with or without 4 g/L acetic acid, under the same conditions described above.

Samples were taken to measure OD and for subsequent high-performance liquid chromatography (HPLC) analysis.

### Analytical procedures

Quantification of glucose, xylose, xylitol, glycerol, acetic acid, formic acid, ethanol, HMF, and furfural was performed by High-Performance Liquid Chromatography (HPLC) using the Alliance HT chromatograph (Waters) with a refractive index detector (Waters 2414) and diode array (Waters 2998). Samples were analysed using an Aminex HPX87H ion exclusion column (300 mm × 7.8 mm, BioRad®), heated at 35°C, with 5 mM H_2_SO_4_ as the mobile phase at a flow rate of 0.6 mL/min. A standard curve with known concentrations of the compounds of interest was analysed by the same procedure. The concentrations of compounds in the samples were determined by comparing the areas of chromatographic peaks obtained with the calibration curves.

### Whole-genome sequencing and analysis of CNVs and SNPs

The genomic DNA sequencing of the AceY.14 strain was performed using the Illumina/HiSEQ 2500 platform at the University of North Carolina sequencing facility (UNC, USA). The SPAdes software (Bankevich et al., 2012) was used for genome assembly. For mutation identification, the reads generated from the AceY.14 strain were aligned with the assembly of the parental strain LVY34.4 using the Bowtie2 aligner (Langmead and Salzberg, 2012), configured to allow mismatches and indels. The identification of single nucleotide polymorphisms (SNPs) was performed by combining the Freebayes (Garrison and Marth, 2012) and GATK package (DePristo et al., 2011) programs. A custom PERL script was used to identify polymorphisms that arose in AceY.14 compared to the parental strain, discarding SNPs identified as assembly errors. Copy number variation (CNV) analysis in strain AceY.14 was performed using the cn.MOPS program (Klambauer et al., 2012), using the results of read alignments from AceY.14 against the genome of the parental strain LVY34.4.

### Yeast transformations and CRISPR/Cas system

For all *S. cerevisiae* transformations, the lithium acetate method with the addition of DMSO prior to heat shock was used (Gietz and Schiestl, 2007; Hill et al., 1991).

Individual knockout mutants of *ZWF1* and *CLN3* genes in the C5TY strain were constructed through homologous recombination with hphMX4 deletion cassette amplified from the plasmid pAG32 (Goldstein and McCusker, 1999) using primers with 42 bp-homology regions up- and downstream the target genes. Double mutants were constructed by deleting the *ISU1* gene through homologous recombination using the KanMX4 cassette (Wach et al., 1994), amplified from the pFA6KanMX4 plasmid. Single and double mutants were selected on media supplemented with 200 mg/L of hygromycin or geneticin.

Mutant cells with point mutations were constructed using the CRISPR/Cas9 EasyGuide system (Jacobus et al., 2022). The single deletion strains (*zwf1*Δ*::hphMX4* and *cln3*Δ*::hphMX4*) were used as the background and transformed with the cas9 plasmid pJACas-K. The target gene sequences with the point mutations were amplified from the evolved parental strain AceY.14 and used as DNA donors in a second transformation, where the donors and the pJASPR6 plasmid with the guide RNA (gRNA) designed to target the hphMX4 sequence, aiming the replacement of the resistance gene by the donor with the mutated sequence.

All transformants were confirmed by PCR and the insertion of the point mutations was confirmed by Sanger sequencing. All primers used in this study are listed in the Supplementary Table S4.

### Spot assay in acetic acid

Cells were grown overnight in 20 mL of YPD medium at 30°C and 200 rpm, centrifuged, washed, and suspended in sterile water at an initial OD_600_ of 1. Serial dilutions were prepared in microtiter plates for 10^-1^, 10^-2^,10^-3^ and 10^-4^ concentrations. Samples (5 μl) of each dilution were inoculated on solid YP medium with 20 g/L of xylose supplemented with 4, 4.5, or 5 g/L of acetic acid, without pH adjustment. All experiments were made in triplicate, and plates were incubated at 30°C for 4 days. The growth in the medium without inhibitor was used as a control. The results were analyzed in the ImageJ software as previously described for cellular growth quantification (Petropavlovskiy et al., 2020).

### Determination of NADP^+^/NADPH ratio

An overnight culture of cells grown in YPD medium was washed and inoculated in triplicate into YP medium containing 5% xylose, with a starting ODLLL of 1. Cultures were incubated at 30 °C with shaking at 200 rpm until reaching mid-log phase (ODLLL = 6– 8). Cells were then harvested by centrifugation at 4000 rpm for 2 minutes and resuspended in PBS buffer to an equivalent cell density (ODLLL = 1). Samples were prepared for individual measurement of NADPL and NADPH levels using the NADP/NADPH-Glo™ Assay (Promega, Cat. No. G9081), following the manufacturer’s protocol. Reactions were carried out in a 96-well black microtiter plate and incubated at room temperature for 30 minutes. Luminescence was measured using a SpectraMax® M3 luminometer. Data were normalized to a buffer-only control (blank), and the NADPL/NADPH ratio was subsequently calculated.

### Diploidization of AceY.14

The AceY.14 strain was transformed with the plasmid pSHO (personal communication), containing the gene encoding the endonuclease HO instead of the gene encoding the recombinase Cre in the pSH65 plasmid (Gueldener, 2002). After selection, a transformant was inoculated in YPG medium for eight hours to induce HO by the GAL promoter, and colonies were isolated for diploid screening. Confirmation of the diploid *MAT*α*/MATa* was performed by PCR according (Illuxley et al., 1990).

### Lignocellulosic hydrolysate preparation

Lignocellulosic hydrolysates used in this study were obtained from eucalyptus chips, sugarcane straw and bagasse, and energy cane. The hydrolysates were prepared at the GranBio/Celere laboratories and kindly provided by the company for the study. One hundred and fifty grams of each material were pretreated using steam explosion, maintaining a temperature of 183°C for five minutes before the explosion. The pretreated material was hydrolyzed with a total solids loading of 15% (w/v) using the Novozymes CTEC3 commercial cocktail, with a pH of 5 at 50°C for 72 hours. Samples were collected for quantification of released sugars and inhibitors (Table S2). For fermentation, urea was added at a final concentration of 1 g/L, and the final pH was adjusted to 5.5 with KOH. No other components were added for fermentation.

## Acknowledgements

We are thankful to Fundação de Amparo à Pesquisa do Estado de São Paulo (FAPESP, Grant Numbers: 2017/08519-6, 2020/07918-7, 2020/02936-7 and 2019/06942-4) and Serrapilheira Institute (Grant Number: Serra-1708-16205). This study was financed in part by the Coordenação de Aperfeiçoamento de Pessoal de Nível Superior-Brasil (CAPES)—Finance Code 001. We thank Thamy L R Correa for her scientific and artistic contributions on the structural model in Figure 3.

## Funding

This work was supported by Fundação de Amparo à Pesquisa do Estado de São Paulo (FAPESP, Grant Numbers: 2017/08519-6, 2020/07918-7, 2020/02936-7 and 2019/06942-4) and Serrapilheira Institute (Grant Number: Serra-1708-16205). This study was financed in part by the Coordenação de Aperfeiçoamento de Pessoal de Nível Superior-Brasil (CAPES)—Finance Code 001.

## Contributions

**Leandro Vieira dos Santos:** Conceptualization, Investigation, Methodology, Writing - Original Draft, Writing - Review & Editing, Supervision, Funding acquisition. **Gisele Cristina de Lima Palermo:** Investigation, Writing - Original Draft, Writing - Review & Editing, Visualization, Formal analysis. **Paulo Emílio dos Santos Costa:** Investigation, Writing - Original Draft**. Ludimila Dias Almeida:** Investigation. **Marcelo Falsarella Carazzolle:** Data Curation. **Gonçalo Amarante Guimarães Pereira:** Supervision, Funding acquisition

## Competing interests

The authors declare no competing financial interests.

## Notes

### Competing Interest Statement

The authors have declared no competing interest.

## References

Adler, S.O., Spiesser, T.W., Uschner, F., Münzner, U., Hahn, J., Krantz, M., Klipp, E., 2022. A yeast cell cycle model integrating stress, signaling, and physiology. FEMS Yeast Research 22, foac026. 10.1093/femsyr/foac026

Almeida, J.R., Modig, T., Petersson, A., HähnlHägerdal, B., Lidén, G., GorwalGrauslund, M.F., 2007. Increased tolerance and conversion of inhibitors in lignocellulosic hydrolysates by *Saccharomyces cerevisiae*. J of Chemical Tech & Biotech 82, 340–349. 10.1002/jctb.1676

Antar, M., Lyu, D., Nazari, M., Shah, A., Zhou, X., Smith, D.L., 2021. Biomass for a sustainable bioeconomy: An overview of world biomass production and utilization. Renewable and Sustainable Energy Reviews 139, 110691. 10.1016/j.rser.2020.110691

Bankevich, A., Nurk, S., Antipov, D., Gurevich, A.A., Dvorkin, M., Kulikov, A.S., Lesin, V.M., Nikolenko, S.I., Pham, S., Prjibelski, A.D., Pyshkin, A.V., Sirotkin, A.V., Vyahhi, N., Tesler, G., Alekseyev, M.A., Pevzner, P.A., 2012. SPAdes: A New Genome Assembly Algorithm and Its Applications to Single-Cell Sequencing. Journal of Computational Biology 19, 455–477. 10.1089/cmb.2012.0021

Basso, L.C., De Amorim, H.V., De Oliveira, A.J., Lopes, M.L., 2008. Yeast selection for fuel ethanol production in Brazil. FEMS Yeast Research 8, 1155–1163. 10.1111/j.1567-1364.2008.00428.x

Bellissimi, E., Van Dijken, J.P., Pronk, J.T., Van Maris, A.J.A., 2009. Effects of acetic acid on the kinetics of xylose fermentation by an engineered, xylose-isomerase-based *Saccharomyces cerevisiae* strain. FEMS Yeast Research 9, 358–364. 10.1111/j.1567-1364.2009.00487.x

Benaroudj, N., Lee, D.H., Goldberg, A.L., 2001. Trehalose Accumulation during Cellular Stress Protects Cells and Cellular Proteins from Damage by Oxygen Radicals. Journal of Biological Chemistry 276, 24261–24267. 10.1074/jbc.M101487200

Blank, L.M., Kuepfer, L., Sauer, U., 2005. Large-scale 13C-flux analysis reveals mechanistic principles of metabolic network robustness to null mutations in yeast. Genome Biol 6, R49. 10.1186/gb-2005-6-6-r49

Celton, M., Goelzer, A., Camarasa, C., Fromion, V., Dequin, S., 2012. A constraint-based model analysis of the metabolic consequences of increased NADPH oxidation in *Saccharomyces cerevisiae*. Metabolic Engineering 14, 366–379. 10.1016/j.ymben.2012.03.008

Chen, A., Gibney, P.A., 2022. Intracellular trehalose accumulation via the Agt1 transporter promotes freeze–thaw tolerance in *Saccharomyces cerevisiae*. J Appl Microbiol. 133, 2390–2402. 10.1111/jam.15700

Costanzo, M., Nishikawa, J.L., Tang, X., Millman, J.S., Schub, O., Breitkreuz, K., Dewar, D., Rupes, I., Andrews, B., Tyers, M., 2004. CDK Activity Antagonizes Whi5, an Inhibitor of G1/S Transcription in Yeast. Cell 117, 899–913. 10.1016/j.cell.2004.05.024

Cross, F.R., Blake, C.M., 1993. The Yeast Cln3 Protein Is an Unstable Activator of Cdc28. MOL. CELL. BIOL. 13, 3266–3271. 10.1128/mcb.13.6.3266-3271.1993

De Bruin, R.A.M., Kalashnikova, T.I., Chahwan, C., McDonald, W.H., Wohlschlegel, J., Yates, J., Russell, P., Wittenberg, C., 2006. Constraining G1-Specific Transcription to Late G1 Phase: The MBF-Associated Corepressor Nrm1 Acts via Negative Feedback. Molecular Cell 23, 483–496. 10.1016/j.molcel.2006.06.025

De Bruin, R.A.M., McDonald, W.H., Kalashnikova, T.I., Yates, J., Wittenberg, C., 2004. Cln3 Activates G1-Specific Transcription via Phosphorylation of the SBF Bound Repressor Whi5. Cell 117, 887–898. 10.1016/j.cell.2004.05.025

Demeke, M.M., Dietz, H., Li, Y., Foulquié-Moreno, M.R., Mutturi, S., Deprez, S., Den Abt, T., Bonini, B.M., Liden, G., Dumortier, F., Verplaetse, A., Boles, E., Thevelein, J.M., 2013. Development of a D-xylose fermenting and inhibitor tolerant industrial *Saccharomyces cerevisiae* strain with high performance in lignocellulose hydrolysates using metabolic and evolutionary engineering. Biotechnol Biofuels 6, 89. 10.1186/1754-6834-6-89

DePristo, M.A., Banks, E., Poplin, R., Garimella, K.V., Maguire, J.R., Hartl, C., Philippakis, A.A., Del Angel, G., Rivas, M.A., Hanna, M., McKenna, A., Fennell, T.J., Kernytsky, A.M., Sivachenko, A.Y., Cibulskis, K., Gabriel, S.B., Altshuler, D., Daly, M.J., 2011. A framework for variation discovery and genotyping using next-generation DNA sequencing data. Nat Genet 43, 491–498. 10.1038/ng.806

Diao, L., Liu, Y., Qian, F., Yang, J., Jiang, Y., Yang, S., 2013. Construction of fast xylose-fermenting yeast based on industrial ethanol-producing diploid *Saccharomyces cerevisiae* by rational design and adaptive evolution. BMC Biotechnol 13, 110. 10.1186/1472-6750-13-110

Ding, J., Bierma, J., Smith, M.R., Poliner, E., Wolfe, C., Hadduck, A.N., Zara, S., Jirikovic, M., Van Zee, K., Penner, M.H., Patton-Vogt, J., Bakalinsky, A.T., 2013. Acetic acid inhibits nutrient uptake in *Saccharomyces cerevisiae*: auxotrophy confounds the use of yeast deletion libraries for strain improvement. Appl Microbiol Biotechnol 97, 7405–7416. 10.1007/s00253-013-5071-y

Dos Santos, L.V., De Barros Grassi, M.C., Gallardo, J.C.M., Pirolla, R.A.S., Calderón, L.L., De Carvalho-Netto, O.V., Parreiras, L.S., Camargo, E.L.O., Drezza, A.L., Missawa, S.K., Teixeira, G.S., Lunardi, I., Bressiani, J., Pereira, G.A.G., 2016b. Second-Generation Ethanol: The Need is Becoming a Reality. Industrial Biotechnology 12, 40–57. 10.1089/ind.2015.0017

Dos Santos, L.V., Carazzolle, M.F., Nagamatsu, S.T., Sampaio, N.M.V., Almeida, L.D., Pirolla, R.A.S., Borelli, G., Corrêa, T.L.R., Argueso, J.L., Pereira, G.A.G., 2016a. Unraveling the genetic basis of xylose consumption in engineered *Saccharomyces cerevisiae* strains. Sci Rep 6, 38676. 10.1038/srep38676

Dos Santos, L.V., Neitzel, T., Lima, C.S., De Carvalho, L.M., De Lima, T.B., Ienczak, J.L., Corrêa, T.L.R., Pereira, G.A.G., 2025. Engineering cellular redox homeostasis to optimize ethanol production in xylose-fermenting *Saccharomyces cerevisiae* strains. Microbiological Research 290, 127955. 10.1016/j.micres.2024.127955

Eleutherio, E., Panek, A., De Mesquita, J.F., Trevisol, E., Magalhães, R., 2015. Revisiting yeast trehalose metabolism. Curr Genet 61, 263–274. 10.1007/s00294-014-0450-1

Favaro, L., Jansen, T., Van Zyl, W.H., 2019. Exploring industrial and natural *Saccharomyces cerevisiae* strains for the bio-based economy from biomass: the case of bioethanol. Critical Reviews in Biotechnology 39, 800–816. 10.1080/07388551.2019.1619157

Fernandes, A.R., Mira, N.P., Vargas, R.C., Canelhas, I., Sá-Correia, I., 2005. *Saccharomyces cerevisiae* adaptation to weak acids involves the transcription factor Haa1p and Haa1p- regulated genes. Biochemical and Biophysical Research Communications 337, 95–103. 10.1016/j.bbrc.2005.09.010

Gancedo, J.M., 1998. Yeast Carbon Catabolite Repression. Microbiol Mol Biol Rev 62, 334–361. 10.1128/MMBR.62.2.334-361.1998

Gao, X., Lin, S.-H., Ren, F., Li, J.-T., Chen, J.-J., Yao, C.-B., Yang, H.-B., Jiang, S.-X., Yan, G.-Q., Wang, D., Wang, Y., Liu, Y., Cai, Z., Xu, Y.-Y., Chen, J., Yu, W., Yang, P.-Y., Lei, Q.-Y., 2016. Acetate functions as an epigenetic metabolite to promote lipid synthesis under hypoxia. Nat Commun 7, 11960. 10.1038/ncomms11960

Garland, S.A., Hoff, K., Vickery, L.E., Culotta, V.C., 1999. *Saccharomyces cerevisiae* ISU1 and ISU2: members of a well-conserved gene family for iron-sulfur cluster assembly. Journal of Molecular Biology 294, 897–907. 10.1006/jmbi.1999.3294

Garrison, E., Marth, G., 2012. Haplotype-based variant detection from short-read sequencing. 10.48550/arXiv.1207.3907

Gietz, R.D., Schiestl, R.H., 2007. Large-scale high-efficiency yeast transformation using the LiAc/SS carrier DNA/PEG method. Nat Protoc 2, 38–41. 10.1038/nprot.2007.15

Glick, B.R., 1995. Metabolic load and heterologous gene expression. Biotechnology Advances 13, 247–261. 10.1016/0734-9750(95)00004-A

Goldstein, A.L., McCusker, J.H., 1999. Three new dominant drug resistance cassettes for gene disruption in *Saccharomyces cerevisiae*. Yeast 15, 1541–1553. 10.1002/(SICI)1097-0061(199910)15:14<1541::AID-YEA476>3.0.CO;2-K

Gong, X., Zhang, J., Gan, Q., Teng, Y., Hou, J., Lyu, Y., Liu, Z., Wu, Z., Dai, R., Zou, Y., Wang, X., Zhu, D., Zhu, H., Liu, T., Yan, Y., 2024. Advancing microbial production through artificial intelligence-aided biology. Biotechnology Advances 74, 108399. 10.1016/j.biotechadv.2024.108399

Gorsich, S.W., Dien, B.S., Nichols, N.N., Slininger, P.J., Liu, Z.L., Skory, C.D., 2006. Tolerance to furfural-induced stress is associated with pentose phosphate pathway genes ZWF1, GND1, RPE1, and TKL1 in *Saccharomyces cerevisiae*. Appl Microbiol Biotechnol 71, 339–349. 10.1007/s00253-005-0142-3

Gueldener, U., 2002. A second set of loxP marker cassettes for Cre-mediated multiple gene knockouts in budding yeast. Nucleic Acids Research 30, 23e–223. 10.1093/nar/30.6.e23

Hao, H., Yao, M., Wang, Y., Zhang, C., Liu, Z., Nielsen, J., Shi, S., Xiao, W., Yuan, Y., 2024. Extending the G1 phase improves the production of lipophilic compounds in yeast by boosting enzyme expression and increasing cell size. PNAS 121, e2413486121. 10.1073/pnas.2413486121

Hill, J., Donald, K.A.I.G., Griffiths, D.E., 1991. DMSO-enhanced whole cell yeast transformation. Nucl Acids Res 19, 5791–5791. 10.1093/nar/19.20.5791

Illuxley, C., Green, E.D., Dunbam, I., 1990. Rapid assessment of *S. cerevisiae* mating type by PCR. Trends in Genetics 6, 236. 10.1016/0168-9525(90)90190-H

Jacobus, A.P., Barreto, J.A., De Bem, L.S., Menegon, Y.A., Fier, Í., Bueno, J.G.R., Dos Santos, L.V., Gross, J., 2022. EasyGuide Plasmids Support in Vivo Assembly of gRNAs for CRISPR/Cas9 Applications in *Saccharomyces cerevisiae*. ACS Synth. Biol. 11, 3886–3891. 10.1021/acssynbio.2c00348

Jacobus, A.P., Cavassana, S.D., De Oliveira, I.I., Barreto, J.A., Rohwedder, E., Frazzon, J., Basso, T.P., Basso, L.C., Gross, J., 2024. Optimal trade-off between boosted tolerance and growth fitness during adaptive evolution of yeast to ethanol shocks. Biotechnol Biofuels 17, 63. 10.1186/s13068-024-02503-7

Jang, W.D., Kim, G.B., Kim, Y., Lee, S.Y., 2022. Applications of artificial intelligence to enzyme and pathway design for metabolic engineering. Current Opinion in Biotechnology 73, 101–107. 10.1016/j.copbio.2021.07.024

Jeppsson, M., Johansson, B., Hahn-Hägerdal, B., Gorwa-Grauslund, M.F., 2002. Reduced Oxidative Pentose Phosphate Pathway Flux in Recombinant Xylose-Utilizing *Saccharomyces cerevisiae* Strains Improves the Ethanol Yield from Xylose. Appl Environ Microbiol 68, 1604–1609. 10.1128/AEM.68.4.1604-1609.2002

Jeppsson, Marie, Johansson, B., Jensen, P.R., HahnlHägerdal, B., GorwalGrauslund, M.F., 2003. The level of glucosel6lphosphate dehydrogenase activity strongly influences xylose fermentation and inhibitor sensitivity in recombinant *Saccharomyces cerevisiae* strains. Yeast 20, 1263–1272. 10.1002/yea.1043

Jeppsson, M, Traff, K., Johansson, B., Hahnhagerdal, B., Gorwagrauslund, M., 2003. Effect of enhanced xylose reductase activity on xylose consumption and product distribution in xylose-fermenting recombinant. FEMS Yeast Research 3, 167–175. 10.1016/S1567-1356(02)00186-1

Jönsson, L.J., Martín, C., 2016. Pretreatment of lignocellulose: Formation of inhibitory by-products and strategies for minimizing their effects. Bioresource Technology 199, 103–112. 10.1016/j.biortech.2015.10.009

Karhumaa, K., Sanchez, R.G., Hahn-Hägerdal, B., Gorwa-Grauslund, M.-F., 2007. Comparison of the xylose reductase-xylitol dehydrogenase and the xylose isomerase pathways for xylose fermentation by recombinant *Saccharomyces cerevisiae*. Microb Cell Fact 6, 5. 10.1186/1475-2859-6-5

Klambauer, G., Schwarzbauer, K., Mayr, A., Clevert, D.-A., Mitterecker, A., Bodenhofer, U., Hochreiter, S., 2012. cn.MOPS: mixture of Poissons for discovering copy number variations in next-generation sequencing data with a low false discovery rate. Nucleic Acids Research 40, e69–e69. 10.1093/nar/gks003

Kotaka, M., Gover, S., Vandeputte-Rutten, L., Au, S.W.N., Lam, V.M.S., Adams, M.J., 2005. Structural studies of glucose-6-phosphate and NADP ^+^ binding to human glucose-6-phosphate dehydrogenase. Acta Cryst. 61, 495–504. 10.1107/S0907444905002350

Kuyper, M., Harhangi, H., Stave, A., Winkler, A., Jetten, M., Delaat, W., Denridder, J., Opdencamp, H., Vandijken, J., Pronk, J., 2003. High-level functional expression of a fungal xylose isomerase: the key to efficient ethanolic fermentation of xylose by *Saccharomyces cerevisiae*l? FEMS Yeast Research 4, 69–78. 10.1016/S1567-1356(03)00141-7

Kuyper, M., Toirkens, M., Diderich, J., Winkler, A., Vandijken, J., Pronk, J., 2005. Evolutionary engineering of mixed-sugar utilization by a xylose-fermenting strain. FEMS Yeast Research 5, 925–934. 10.1016/j.femsyr.2005.04.004

Langmead, B., Salzberg, S.L., 2012. Fast gapped-read alignment with Bowtie 2. Nat Methods 9, 357–359. 10.1038/nmeth.1923

Lee, S.-B., Tremaine, M., Place, M., Liu, L., Pier, A., Krause, D.J., Xie, D., Zhang, Y., Landick, R., Gasch, A.P., Hittinger, C.T., Sato, T.K., 2021. Crabtree/Warburg-like aerobic xylose fermentation by engineered *Saccharomyces cerevisiae*. Metabolic Engineering 68, 119–130. 10.1016/j.ymben.2021.09.008

Lee, S.-M., Jellison, T., Alper, H.S., 2014. Systematic and evolutionary engineering of a xylose isomerase-based pathway in *Saccharomyces cerevisiae* for efficient conversion yields. Biotechnol Biofuels 7, 122. 10.1186/s13068-014-0122-x

Lewis Liu, Z., Ma, M., Song, M., 2009. Evolutionarily engineered ethanologenic yeast detoxifies lignocellulosic biomass conversion inhibitors by reprogrammed pathways. Mol Genet Genomics 282, 233–244. 10.1007/s00438-009-0461-7

Lill, R., Hoffmann, B., Molik, S., Pierik, A.J., Rietzschel, N., Stehling, O., Uzarska, M.A., Webert, H., Wilbrecht, C., Mühlenhoff, U., 2012. The role of mitochondria in cellular iron–sulfur protein biogenesis and iron metabolism. Biochimica et Biophysica Acta (BBA) - Molecular Cell Research 1823, 1491–1508. 10.1016/j.bbamcr.2012.05.009

Ling, H., Teo, W., Chen, B., Leong, S.S.J., Chang, M.W., 2014. Microbial tolerance engineering toward biochemical production: from lignocellulose to products. Current Opinion in Biotechnology 29, 99–106. 10.1016/j.copbio.2014.03.005

Madhavan, A., Tamalampudi, S., Ushida, K., Kanai, D., Katahira, S., Srivastava, A., Fukuda, H., Bisaria, V.S., Kondo, A., 2009. Xylose isomerase from polycentric fungus *Orpinomyces*: gene sequencing, cloning, and expression in *Saccharomyces cerevisiae* for bioconversion of xylose to ethanol. Appl Microbiol Biotechnol 82, 1067–1078. 10.1007/s00253-008-1794-6

Mahmud, S.A., Hirasawa, T., Shimizu, H., 2010. Differential importance of trehalose accumulation in *Saccharomyces cerevisiae* in response to various environmental stresses. J. BIOSCI. BIOENG. 109, 262–266. 10.1016/j.jbiosc.2009.08.500

Mao, J., Zhang, H., Chen, Yu, Wei, L., Liu, J., Nielsen, J., Chen, Yun, Xu, N., 2024. Relieving metabolic burden to improve robustness and bioproduction by industrial microorganisms. Biotechnology Advances 74, 108401. 10.1016/j.biotechadv.2024.108401

McMillan, J.D., 1993. Xylose fermentation to ethanol. A review (No. NREL/TP--421-4944, 10117941). 10.2172/10117941

Mira, N.P., Teixeira, M.C., Sá-Correia, I., 2010. Adaptive Response and Tolerance to Weak Acids in *Saccharomyces cerevisiae*l: A Genome-Wide View. OMICS: A Journal of Integrative Biology 14, 525–540. 10.1089/omi.2010.0072

Myers, K.S., Riley, N.M., MacGilvray, M.E., Sato, T.K., McGee, M., Heilberger, J., Coon, J.J., Gasch, A.P., 2019. Rewired cellular signaling coordinates sugar and hypoxic responses for anaerobic xylose fermentation in yeast. PLoS Genet 15, e1008037. 10.1371/journal.pgen.1008037

Nascimento, V.M., Ienczak, J.L., Boni, R., Maciel Filho, R., Rabelo, S.C., 2023. Mapping potential solvents for inhibitors removal from sugarcane bagasse hemicellulosic hydrolysate and its impact on fermentability. Industrial Crops and Products 192, 116023. 10.1016/j.indcrop.2022.116023

Osiro, K.O., Borgström, C., Brink, D.P., Fjölnisdóttir, B.L., Gorwa-Grauslund, M.F., 2019. Exploring the xylose paradox in *Saccharomyces cerevisiae* through in vivo sugar signalomics of targeted deletants. Microb Cell Fact 18, 88. 10.1186/s12934-019-1141-x

Paalman, J.W.G., Verwaal, R., Slofstra, S.H., Verkleij, A.J., Boonstra, J., Verrips, C.T., 2003. Trehalose and glycogen accumulation is related to the duration of the G _1_ phase of *Saccharomyces cerevisiae*. FEMS Yeast Research 3, 261–268. 10.1111/j.1567-1364.2003.tb00168.x

Palermo, G.C.D.L., Coutouné, N., Bueno, J.G.R., Maciel, L.F., Dos Santos, L.V., 2021. Exploring metal ion metabolisms to improve xylose fermentation in *Saccharomyces cerevisiae*. Microbial Biotechnology 14, 2101–2115. 10.1111/1751-7915.13887

Palmqvist, E., Hahn-Hägerdal, B., 2000. Fermentation of lignocellulosic hydrolysates. II: inhibitors and mechanisms of inhibition. Bioresource Technology 74, 25–33. 10.1016/S0960-8524(99)00161-3

Petropavlovskiy, A.A., Tauro, M.G., Lajoie, P., Duennwald, M.L., 2020. A Quantitative Imaging-Based Protocol for Yeast Growth and Survival on Agar Plates. STAR Protocols 1, 100182. 10.1016/j.xpro.2020.100182

Piotrowski, J.S., Zhang, Y., Bates, D.M., Keating, D.H., Sato, T.K., Ong, I.M., Landick, R., 2014. Death by a thousand cuts: the challenges and diverse landscape of lignocellulosic hydrolysate inhibitors. Front. Microbiol. 5. 10.3389/fmicb.2014.00090

Qi, X., Zha, J., Liu, G.-G., Zhang, W., Li, B.-Z., Yuan, Y.-J., 2015. Heterologous xylose isomerase pathway and evolutionary engineering improve xylose utilization in *Saccharomyces cerevisiae*. Front. Microbiol. 6. 10.3389/fmicb.2015.01165

Qiu, Y., Liu, Wei, Wu, M., Bao, H., Sun, X., Dou, Q., Jia, H., Liu, Weifeng, Shen, Y., 2024. Construction of an alternative NADPH regeneration pathway improves ethanol production in *Saccharomyces cerevisiae* with xylose metabolic pathway. Synthetic and Systems Biotechnology 9, 269–276. 10.1016/j.synbio.2024.02.004

Roque, L.R., Morgado, G.P., Nascimento, V.M., Ienczak, J.L., Rabelo, S.C., 2019. Liquid-liquid extraction: A promising alternative for inhibitors removing of pentoses fermentation. Fuel 242, 775–787. 10.1016/j.fuel.2018.12.130

Salas-Navarrete, P.C., De Oca Miranda, A.I.M., Martínez, A., Caspeta, L., 2021. Evolutionary and reverse engineering to increase *Saccharomyces cerevisiae* tolerance to acetic acid, acidic pH, and high temperature. Appl Microbiol Biotechnol 106, 383–399. 10.1007/s00253-021-11730-z

Sànchez I Nogué, V., Narayanan, V., Gorwa-Grauslund, M.F., 2013. Short-term adaptation improves the fermentation performance of *Saccharomyces cerevisiae* in the presence of acetic acid at low pH. Appl Microbiol Biotechnol 97, 7517–7525. 10.1007/s00253-013-5093-5

Sato, T.K., Tremaine, M., Parreiras, L.S., Hebert, A.S., Myers, K.S., Higbee, A.J., Sardi, M., McIlwain, S.J., Ong, I.M., Breuer, R.J., Avanasi Narasimhan, R., McGee, M.A., Dickinson, Q., La Reau, A., Xie, D., Tian, M., Reed, J.L., Zhang, Y., Coon, J.J., Hittinger, C.T., Gasch, A.P., Landick, R., 2016. Directed Evolution Reveals Unexpected Epistatic Interactions That Alter Metabolic Regulation and Enable Anaerobic Xylose Use by *Saccharomyces cerevisiae*. PLoS Genet 12, e1006372. 10.1371/journal.pgen.1006372

Shannon, P., Markiel, A., Ozier, O., Baliga, N.S., Wang, J.T., Ramage, D., Amin, N., Schwikowski, B., Ideker, T., 2003. Cytoscape: A Software Environment for Integrated Models of Biomolecular Interaction Networks. Genome Res. 13, 2498–2504. 10.1101/gr.1239303

Stratford, M., Anslow, P.A., 1998. Evidence that sorbic acid does not inhibit yeast as a classic ‘weak acid preservative.’ Letters in Applied Microbiology 27, 203–206. 10.1046/j.1472-765X.1998.00424.x

Sun, Y., Kong, M., Li, X., Li, Q., Xue, Q., Hou, J., Jia, Z., Lei, Z., Xiao, W., Shi, S., Cao, L., 2022. Metabolic and Evolutionary Engineering of Diploid Yeast for the Production of First- and Second-Generation Ethanol. Front. Bioeng. Biotechnol. 9, 835928. 10.3389/fbioe.2021.835928

Talia, S.D., Skotheim, J.M., Bean, J.M., Siggia, E.D., Cross, F.R., 2007. The effects of molecular noise and size control on variability in the budding yeast cell cycle. Nature 448, 947–951. 10.1038/nature06072

Teufel, L., Tummler, K., Flöttmann, M., Herrmann, A., Barkai, N., Klipp, E., 2019. A transcriptome-wide analysis deciphers distinct roles of G1 cyclins in temporal organization of the yeast cell cycle. Sci Rep 9, 3343. 10.1038/s41598-019-39850-7

Travesa, A., Kalashnikova, T.I., De Bruin, R.A.M., Cass, S.R., Chahwan, C., Lee, D.E., Lowndes, N.F., Wittenberg, C., 2013. Repression of G _1_ /S Transcription Is Mediated via Interaction of the GTB Motifs of Nrm1 and Whi5 with Swi6. Molecular and Cellular Biology 33, 1476– 1486. 10.1128/MCB.01333-12

Tyers, M., Tokiwa, G., Nash, R., Futcher, B., 1992. The Cln3-Cdc28 kinase complex of *S. cerevisiae* is regulated by proteolysis and phosphorylation. The EMBO Journal 11, 1773–1784. 10.1002/j.1460-2075.1992.tb05229.x

Ullah, A., Orij, R., Brul, S., Smits, G.J., 2012. Quantitative Analysis of the Modes of Growth Inhibition by Weak Organic Acids in *Saccharomyces cerevisiae*. Appl Environ Microbiol 78, 8377–8387. 10.1128/AEM.02126-12

Verduyn, C., Postma, E., Scheffers, W.A., Van Dijken, J.P., 1992. Effect of benzoic acid on metabolic fluxes in yeasts: A continuouslculture study on the regulation of respiration and alcoholic fermentation. Yeast 8, 501–517. 10.1002/yea.320080703

Verho, R., Londesborough, J., Penttilä, M., Richard, P., 2003. Engineering Redox Cofactor Regeneration for Improved Pentose Fermentation in *Saccharomyces cerevisiae*. Appl Environ Microbiol 69, 5892–5897. 10.1128/AEM.69.10.5892-5897.2003

Verhoeven, M.D., Lee, M., Kamoen, L., Van Den Broek, M., Janssen, D.B., Daran, J.-M.G., Van Maris, A.J.A., Pronk, J.T., 2017. Mutations in PMR1 stimulate xylose isomerase activity and anaerobic growth on xylose of engineered *Saccharomyces cerevisiae* by influencing manganese homeostasis. Sci Rep 7, 46155. 10.1038/srep46155

Wach, A., Brachat, A., Pöhlmann, R., Philippsen, P., 1994. New heterologous modules for classical or PCRlbased gene disruptions in *Saccharomyces cerevisiae*. Yeast 10, 1793–1808. 10.1002/yea.320101310

Wagner, E.R., Myers, K.S., Riley, N.M., Coon, J.J., Gasch, A.P., 2019. PKA and HOG signaling contribute separable roles to anaerobic xylose fermentation in yeast engineered for biofuel production. PLoS ONE 14, e0212389. 10.1371/journal.pone.0212389

Wagner, E.R., Nightingale, N.M., Jen, A., Overmyer, K.A., McGee, M., Coon, J.J., Gasch, A.P., 2023. PKA regulatory subunit Bcy1 couples growth, lipid metabolism, and fermentation during anaerobic xylose growth in *Saccharomyces cerevisiae*. PLoS Genet 19, e1010593. 10.1371/journal.pgen.1010593

Wei, S., Liu, Y., Wu, M., Ma, T., Bai, X., Hou, J., Shen, Y., Bao, X., 2018. Disruption of the transcription factors Thi2p and Nrm1p alleviates the post-glucose effect on xylose utilization in *Saccharomyces cerevisiae*. Biotechnol Biofuels 11, 112. 10.1186/s13068-018-1112-1

Wright, J., Bellissimi, E., De Hulster, E., Wagner, A., Pronk, J.T., Van Maris, A.J.A., 2011. Batch and continuous culture-based selection strategies for acetic acid tolerance in xylose-fermenting *Saccharomyces cerevisiae*: Selection strategies for acetic acid tolerance. FEMS Yeast Research 11, 299–306. 10.1111/j.1567-1364.2011.00719.x

Xu, X., Lv, X., Bi, X., Chen, J., Liu, L., 2024. Genetic circuits for metabolic flux optimization. Trends in Microbiology 32, 791–806. 10.1016/j.tim.2024.01.004

Yaglom, J., Linskens, M.H.K., Sadis, S., Rubin, D.M., Futcher, B., Finley, D., 1995. p34 ^Cdc28^ -Mediated Control of Cln3 Cyclin Degradation. MOL. CELL. BIOL. 15, 731–741. 10.1128/MCB.15.2.731

Yamanaka, K., 1969. Inhibition of d-xylose isomerase by pentitols and d-lyxose. Archives of Biochemistry and Biophysics 131, 502–506. 10.1016/0003-9861(69)90422-6

Yoshiyama, Y., Tanaka, K., Yoshiyama, K., Hibi, M., Ogawa, J., Shima, J., 2015. Trehalose accumulation enhances tolerance of *Saccharomyces cerevisiae* to acetic acid. J. BIOSCI. BIOENG. 119, 172–175. 10.1016/j.jbiosc.2014.06.021

Young, R., Haines, M., Storch, M., Freemont, P.S., 2021. Combinatorial metabolic pathway assembly approaches and toolkits for modular assembly. Metabolic Engineering 63, 81–101. 10.1016/j.ymben.2020.12.001

Yu, J., Li, C., Cheng, Y., Guo, S., Lu, H., Xie, X., Ji, H., Qiao, Y., 2025. Mechanism and improvement of yeast tolerance to biomass-derived inhibitors: A review. Biotechnology Advances 81, 108562. 10.1016/j.biotechadv.2025.108562

Zhang, J., Schneider, C., Ottmers, L., Rodriguez, R., Day, A., Markwardt, J., Schneider, B.L., 2002. Genomic Scale Mutant Hunt Identifies Cell Size Homeostasis Genes in *S. cerevisiae*. Current Biology 12, 1992–2001. 10.1016/S0960-9822(02)01305-2

Zhang, K., Fang, Y.-H., Gao, K.-H., Sui, Y., Zheng, D.-Q., Wu, X.-C., 2017. Effects of genome duplication on phenotypes and industrial applications of *Saccharomyces cerevisiae* strains. Appl Microbiol Biotechnol 101, 5405–5414. 10.1007/s00253-017-8284-7

Zhang, Y.-W., Yang, J.-J., Qian, F.-H., Sutton, K.B., Hjort, C., Wu, W.-P., Jiang, Y., Yang, S., 2024. Engineering a xylose fermenting yeast for lignocellulosic ethanol production. Nat Chem Biol. 10.1038/s41589-024-01771-6

