## Supplementary material for "Epistatic mutations in ISC metabolism synergize with cell cycle regulation and the PPP to enhance xylose fermentation and acetic acid tolerance in industrial yeast"

**Supplementary Figures**

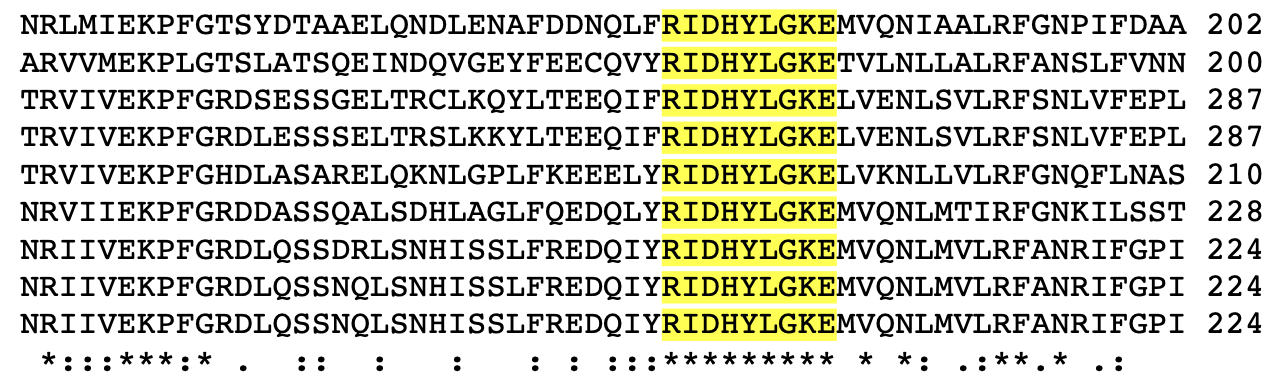

*[Leuconostoc mesenteroides](https://www.uniprot.org/taxonomy/1245" \o "Leuconostoc mesenteroides, taxon ID 1245)* (P11411)

*Escherichia coli*, K12 (P0AC53)

*Arabidopsis thaliana* (Q43727)

*Solanum tuberosum* (Q43839)

*Saccharomyces cerevisiae*, S288C (P11412)

*Drosophila melanogaster* (P12646)

*Homo sapiens* (P11413)

*Rattus norvegicus* (P05370)

*Mus musculus* (Q00612)

**Supplementary Figure S1.** The point mutation E191 of the *ZWF1* gene is in a region with conserved residues between species, as in plants (e.g., *Arabidopsis thaliana* and *Solanum tuberosum*), mammals (e.g., *Homo sapiens* and *Mus musculus*), and microorganisms (e.g., *Escherichia coli* and *Saccharomyces cerevisiae*). Data was generated using the Uniprot align tool.

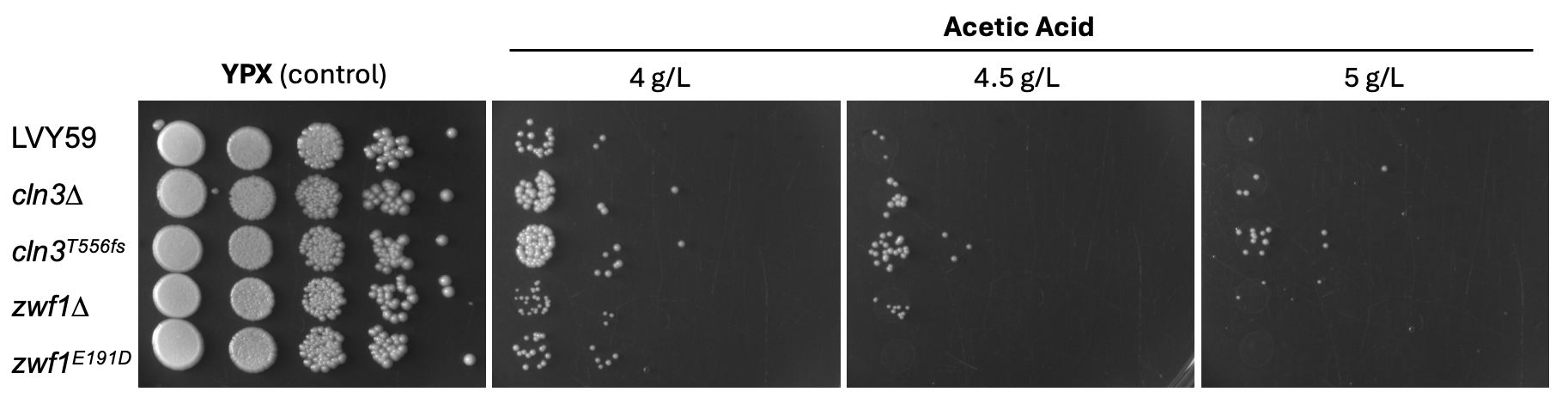

**Supplementary Figure S2.** Spot assay in solid YP medium with 20 g/L of xylose supplemented with 4, 4.5, or 5 g/L of acetic acid, without pH adjustment. Initial OD_600_ of 1 was established before a serial dilution. All experiments were made in triplicate and plates were incubated at 30°C for 4 days. The growth in the medium without inhibitor was used as a control.

**B**

**A**

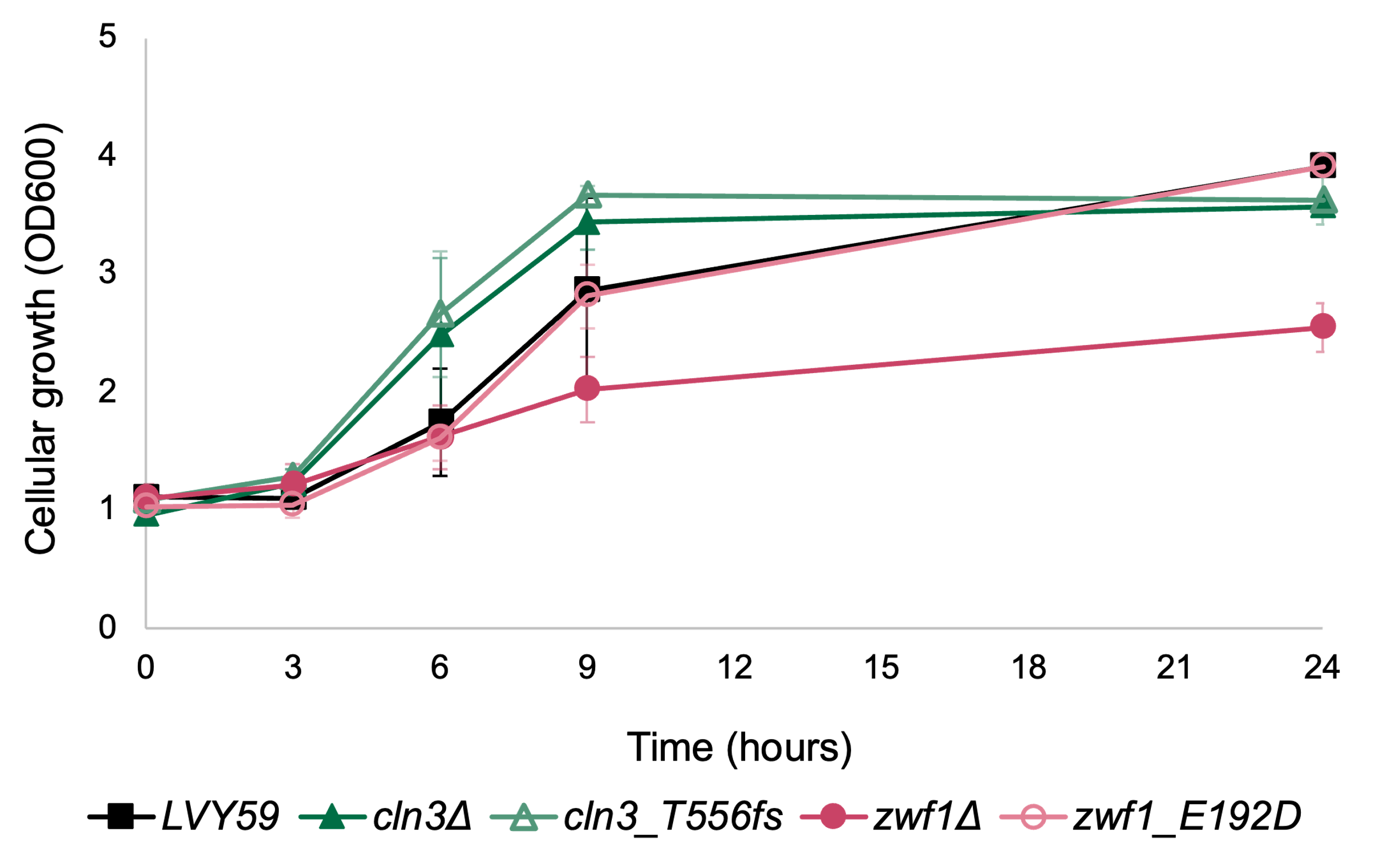

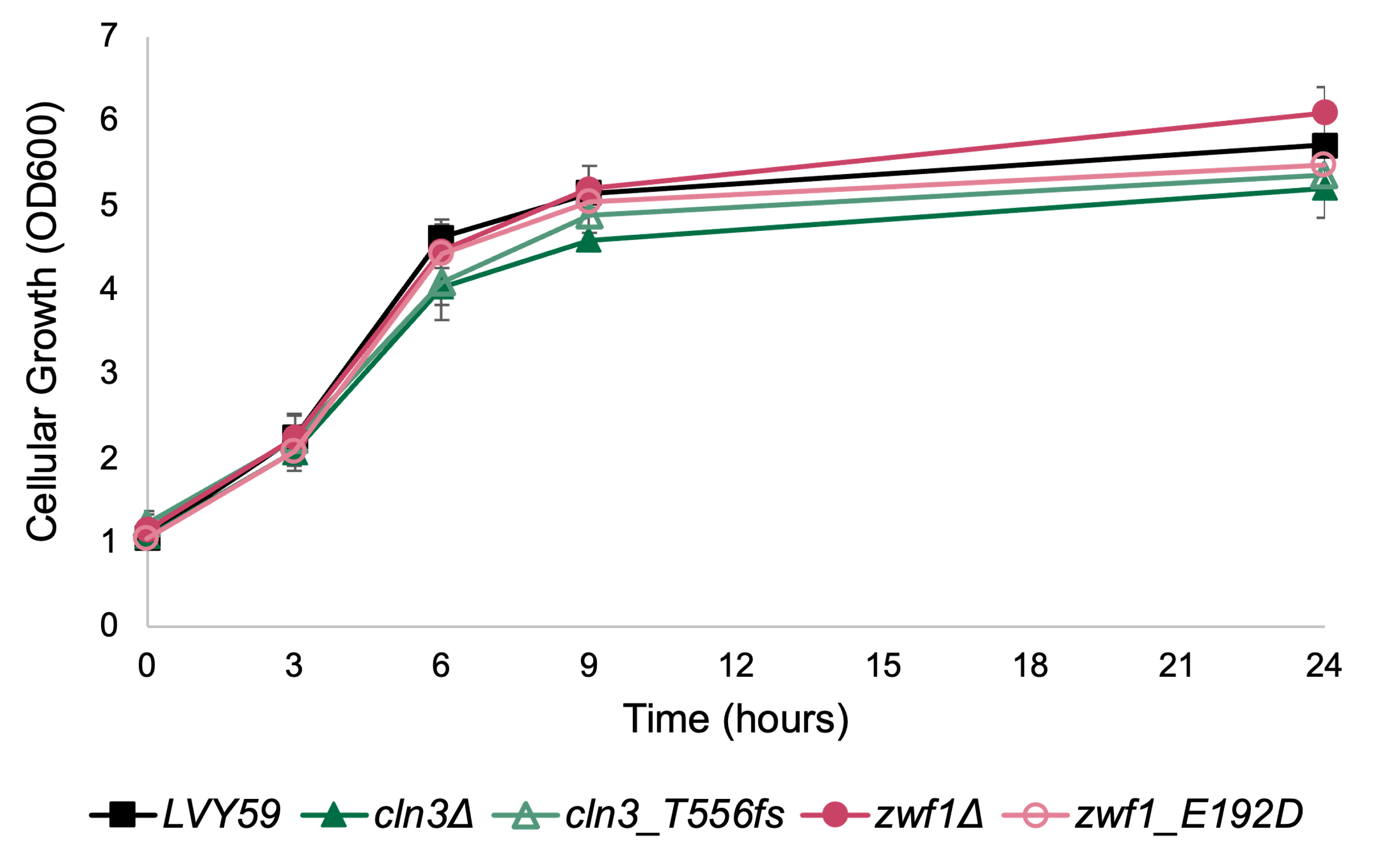

**Supplementary Figure S3.** Cellular growth of *ZWF1* and *CLN3* mutants compared to the control LVY59 in liquid YP media with 20 g/L of glucose, with (A) or without (B) supplementation of 4 g/L of acetic acid, without pH adjustment (pH ~4.3).

Cellular concentration (OD600)

| Time (hours) | LVY59 | cln3∆ | cln3_T556fs | zwf1∆ | zwf1_E192D |
| --- | --- | --- | --- | --- | --- |
| 0 | 1.13125 | 1.08 | 1.106 | 1.082 | 1.16667 |
| 16 | 1.815 | 1.456 | 1.506 | 1.476 | 1.655 |
| 24 | 3.1025 | 2.642 | 2.818 | 5.654 | 4.34833 |
| 32 | 4.25 | 3.606 | 3.54 | 8.808 | 9.50333 |
| 40 | 5.932 | 6.56667 | 6.03333 | 11.036 | 13.06 |
| 48 | 8.69 | 7.308 | 7.874 | 11.148 | 16.6567 |
| 56 | 10.715 | 8.376 | 9.468 | 10.596 | 15.3467 |
| 64 | 11.2 | 9.53333 | 10 | 10.4 |  |
| 72 | 13.6175 | 9.456 | 10.852 | 10.808 | 14.8867 |

**Supplementary Figure S4.** The growth characteristic of *ZWF1* and *CLN3* mutants compared to the control LVY59 in YPX 5%. The error bars represent the standard deviation of biological replicates.

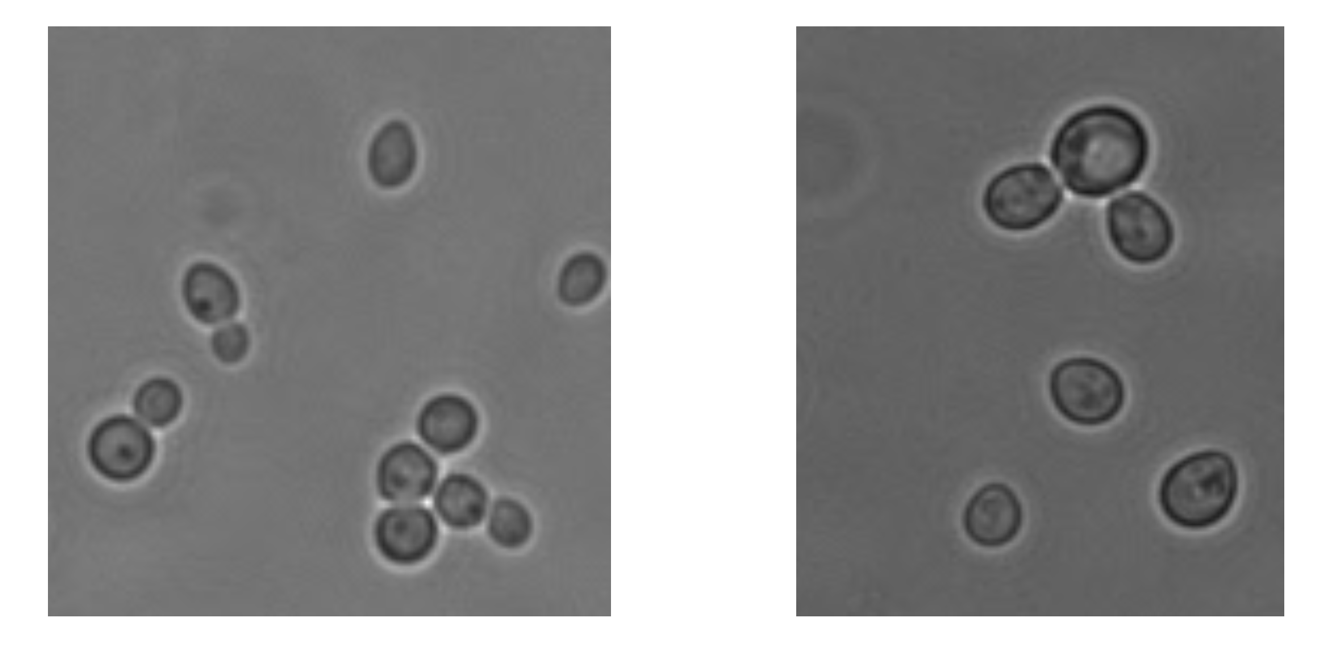

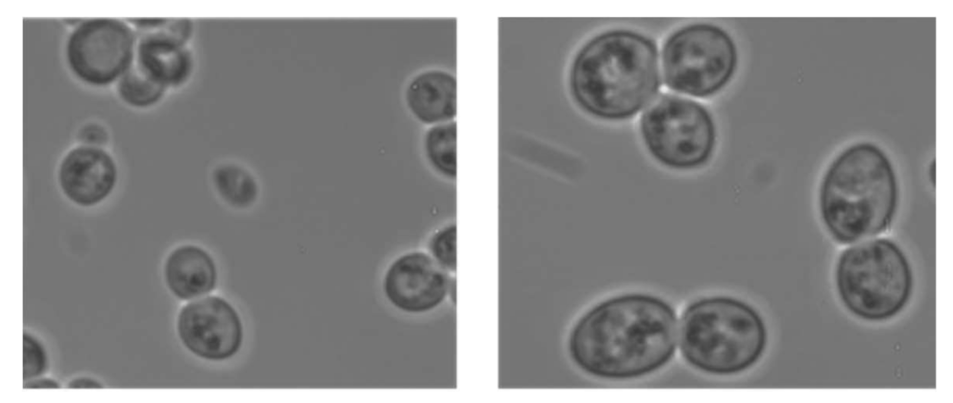

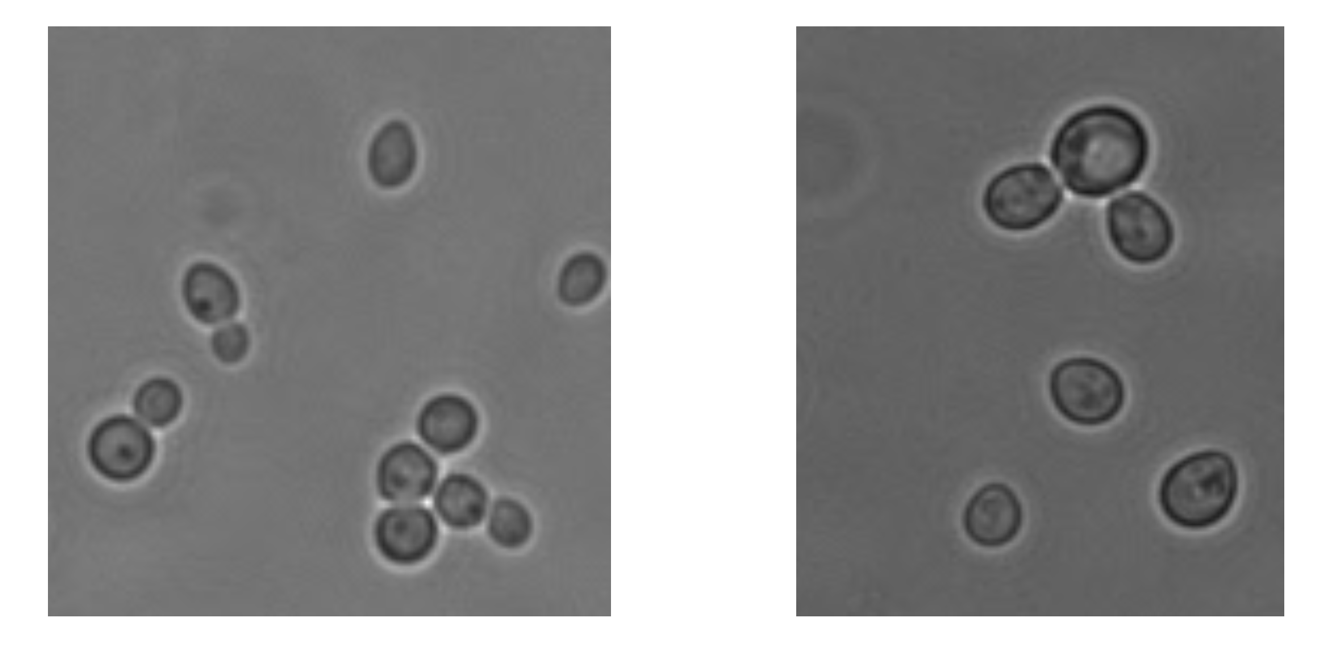

**LVY34.4**

**AceY.14**

**C5TY**

***cln3∆***

**Supplementary Figure S5.** The difference in cell size in AceY.14 is due to *CLN3* mutation. Cells were analyzed in the same magnification compared to their parental strain.

**Supplementary Tables**

**Supplementary Table 1.** Yeast strains used in the study.

| **Strain** | **Relevant genotype/features** | **Reference** |
| --- | --- | --- |
| LVY27 | MATα; CEN5::pTDH1-xylA-tTDH1; gre3Δ; CEN2::pADH1-XKS1-tADH1; CEN8::pADH1-XKS1-tADH1; CEN12::pTDH1-TAL1-tTDH1-pPGK1-RKI1-tPGK1; CEN13::pTDH1-TKL1-tTDH1-pPGK1-RPE1-tPGK1 | Dos Santos et al., 2016 |
| LVY34.4 | LVY27; pOXylATy1 + adaptive evolution in xylose | Dos Santos et al., 2016 |
| C5TY (LVY59) | LVY34.4; ura3Δ; *ISU1 wild type* | Dos Santos et al., 2016 |
| AceY.14 | LVY34.4 + adaptive evolution in acetic acid | This study |
| AceY-2n | AceY.14; MATα/a (diploid) | This study |
| *cln3∆* | LVY59; *cln3∆::hphMX4* | This study |
| *cln3* | cln3∆; *hphMX4::cln3*^T556fs^ | This study |
| *zwf1∆* | LVY59; *cln3∆::hphMX4* | This study |
| *zwf1^E191D^* | *zwf1∆; hphMX4::zwf1^E191D^* | This study |
| *isu1∆* | LVY59; *isu1∆::KanMX4* | This study |
| *isu1∆/cln3∆* | *cnl3∆; isu1∆::KanMX4* | This study |
| *isu1∆/zwf1∆* | *zwf1∆; isu1∆::KanMX4* | This study |

**Supplementary Table 2.** Hydrolysates’ composition.

| **Hydrolysate** | **Initial concentration (g/L)** | | | | | | | | | | | | | | |
| --- | --- | --- | --- | --- | --- | --- | --- | --- | --- | --- | --- | --- | --- | --- | --- |
|  | **Glucose** | | | **Xylose** | | | **Acetic acid** | | | **HMF** | | | **Furfural** | | |
| Sugar cane straw | 55.03 | ± | 1.85 | 32.99 | ± | 1.08 | 4.18 | ± | 0.16 | 0.03 | ± | 0.00 | 0.07 | ± | 0.00 |
| Sugar cane bagasse | 56.09 | ± | 0.31 | 30.26 | ± | 0.15 | 4.53 | ± | 0.20 | 0.41 | ± | 0.02 | 0.04 | ± | 0.00 |
| Energy cane bagasse | 37.73 | ± | 0.07 | 22.72 | ± | 0.05 | 3.24 | ± | 0.04 | 0.20 | ± | 0.00 | 0.02 | ± | 0.00 |
| Eucalyptus | 41.93 | ± | 0.73 | 25.65 | ± | 0.46 | 6.48 | ± | 0.09 | 0.04 | ± | 0.00 | 0.09 | ± | 0.01 |

**Supplementary Table 3.** Fermentation performance of AceY-2 in the different hydrolysates.

| **Hydrolysate** | **Yield (g/g)*** | | | | | | | | | | | **Max. Ethanol volumetric productivity (g/L h^-1^)** | | | |
| --- | --- | --- | --- | --- | --- | --- | --- | --- | --- | --- | --- | --- | --- | --- | --- |
|  | **Ethanol** | | | | **Xylitol** | | | | **Glycerol** | | |  |  |  |  |
| Sugar cane straw | 0.42 | ± | 0.01 | ND | | | | 0.024 | | ± | 0.00 | | 1.574 | ± | 0.050 |
| Sugar cane bagasse | 0.42 | ± | 0.01 | ND | | | | 0.022 | | ± | 0.00 | | 1.856 | ± | 0.043 |
| Energy cane bagasse | 0.40 | ± | 0.01 | ND | | | | 0.021 | | ± | 0.00 | | 1.600 | ± | 0.058 |
| Eucalyptus | 0.41 | ± | 0.01 | 0.02 | | ± | 0.00 | 0.019 | | ± | 0.00 | | 1.466 | ± | 0.040 |

**Supplementary Table 4.** Primers used in this study.

| **Primer Name** | **Sequence (5'-3')** |
| --- | --- |
| ***Gene deletion*** |  |
| Del_ZWF1_F | GAAAGAGTAAATCCAATAGAATAGAAAACCACATAAGGCAAGGCCAGCTGAAGCTTCGTA |
| Del_ZWF1_R | CTAATTATCCTTCGTATCTTCTGGCTTAGTCACGGGCCAAGCGTAAGGTTAGGGAGACCGGCAGATC |
| Del_CLN3_F | TTACTCTCGTTCAAGACACTGATTTGATACGCTTTCTGTACGGCCAGCTGAAGCTTCGTA |
| Del_CLN3_R | TCAGCGAGTTTTCTTGAGGTTGCTACTATCATTAAAATCACAATCCACAGTAGGGAGACCGGCAGATC |
| Del_ISU1_F | ATTGAATAAGGAAAACACAACACATAACACATATTTAACCTGGCCAGCTGAAGCTTCGTA |
| Del_ISU1_R | GAGGGTTTGATCTTGTTCTTGTCCCGGTTATCTTCTTATTCATAGGGAGACCGGCAGATC |
| ***Deletant confirmation*** | |
| Check_ZWF1_F | GATACAAGCTTCCAACGGT |
| Check_ZWF1_R | GCTAATCAAGTGGATAAGACG |
| Check_CLN3_F | TCTGCCAGTCAAATGGATT |
| Check_CLN3_R | ATCAAATATCAGGACGAAGATG |
| Check_ISU1_F | GAGAAGTGGTTCCTTAACCTTAAT |
| Check_ISU1_R | TGATGCCATATACCACACATG |
| ***Sequencing for SNP confirmation*** | |
| ZWF1_int_F | CATAAGGCAAGATGAGTGAA |
